# RNA velocity and protein acceleration from single-cell multiomics experiments

**DOI:** 10.1101/658401

**Authors:** Gennady Gorin, Valentine Svensson, Lior Pachter

**Affiliations:** Division of Chemistry and Chemical Engineering, California Institute of Technology; Division of Biology and Biological Engineering & Department of Computing and Mathematical Sciences, California Institute of Technology

## Abstract

The simultaneous quantification of protein and RNA makes possible the inference of past, present and future cell states from single experimental snapshots. To enable such temporal analysis from multimodal single-cell experiments, we introduce an extension of the RNA velocity method that leverages estimates of unprocessed transcript and protein abundances to extrapolate cell states. We apply the model to four datasets and demonstrate consistency among landscapes and phase portraits.

## Main text

Recent technological innovations that allow for assaying multiple modes of cell states at single-cell resolution are creating opportunities for more detailed biophysical modeling of the molecular biology of cells. Specifically, genome-wide probing of molecular states is revealing detailed information about the functional diversity of cells as determined by gene regulation, transcription, processing, and translation. The ability to probe cell states has been driven by improvements in single-cell RNA sequencing (scRNA-seq) methods^1^ and advances in multiomics^2^. These methods allow researchers to quantify mRNA and protein expression levels in individual cells^3–5^. Furthermore, scRNA-seq can discriminate between nascent and processed transcripts. The recently described *RNA velocity*^6^ method takes advantage of this feature of single-cell RNA-seq to fit a first-order system of ordinary differential equations describing gene-specific splicing^7^ and to infer kinetic trajectories of single cells.

The RNA velocity method uses inferences of abundance of unprocessed transcripts to predict future states of cells with respect to the transcriptional state inferred from the abundance of spliced transcripts. With respect to this reference frame, protein abundances contain information about the abundances of spliced transcripts in the past^8^. To leverage this information, we extend the RNA velocity kinetic model to protein translation, and use single-cell multiomics data with protein quantification to infer, along with the present and future state of cells, the *past* state of each cell in four peripheral blood mononuclear cell (PBMC) datasets (**Fig. 1a**).

**Figure 1.**
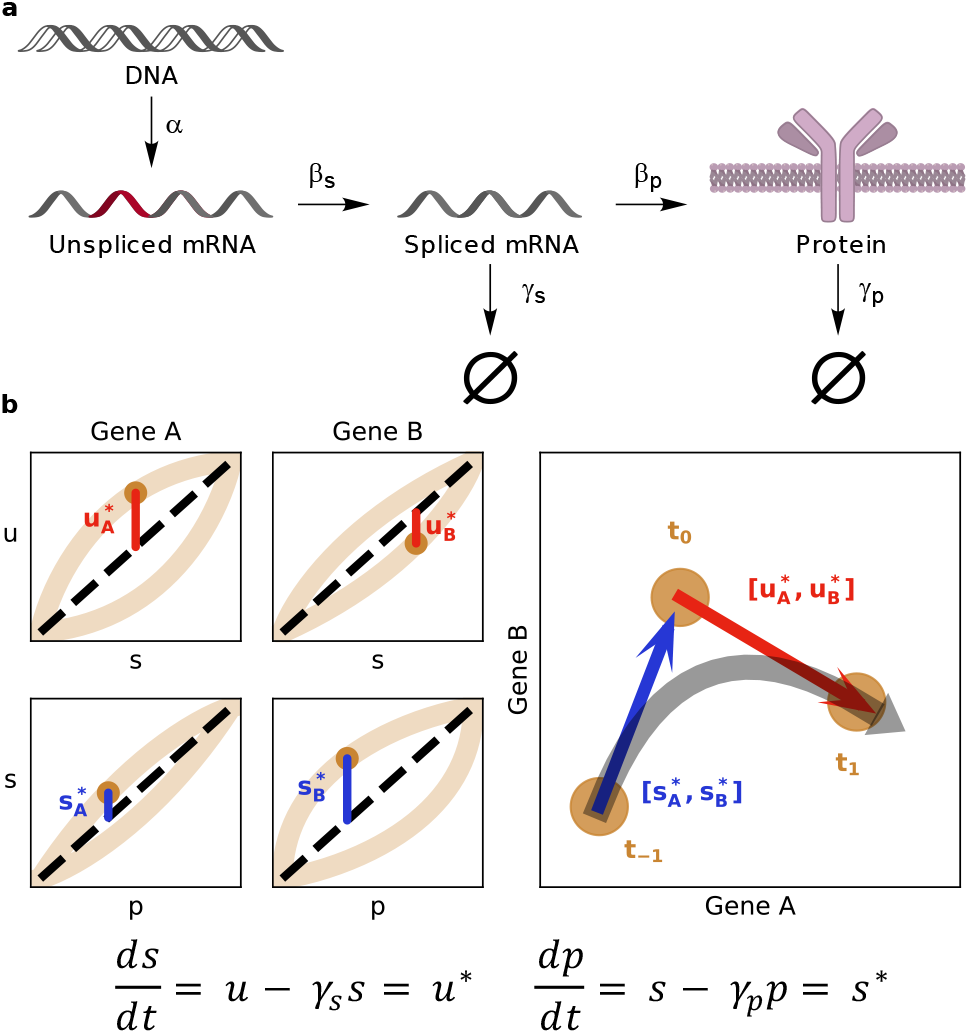
Model structure and parameter inference. (a) A single gene’s information transfer through transcription at rate *α*, splicing at rate *β*_*s*_ · *u*, and translation at rate *β*_*p*_ · *s*. (b) Estimation of equilibrium states *u* = *γ*_*s*_ · *s* and *s* = *γ*_*s*_ · *p* (black, dashed) from imputed gene-specific population data (light brown), followed by calculation of gene- and cell-specific distance from equilibrium (RNA: red, protein: blue) and inference of cell-specific mobility in state space (RNA: red arrow, protein: blue arrow, combined: grey curved arrow).

The net accumulation or depletion of unspliced mRNA, spliced mRNA, and protein, scales with distance from an estimated expression equilibrium (**Fig. 1b**, **Supplementary Note**). Briefly, if a gene’s current (t_0_) unspliced mRNA level is high relative to equilibrium, we infer that the gene is currently upregulated and future (t_1_) spliced mRNA level will be high. Analogously, if the current protein level is high, we infer that the gene is currently downregulated and past (t_−1_) spliced mRNA level was high. Unlike methods that require time-series measurements^9–11^, our method estimates protein translation dynamics from a single time-point.

The approximately linear gene-specific phase plots (**Supplementary Figs. 1-4**) are qualitatively consistent with a first-order model of protein production, although we do observe some deviations by cell type. A subset of high-abundance gene/protein pairs were used to estimate the gene-specific protein accumulation rates (**Supplementary Table 1**). For comparison, the immunoglobulin-coding RNA phase plots (**Supplementary Figs. 5-8**) are quite sparse; for RNA velocity, we used a broad panel of genes with robust unspliced detection. We extrapolated the cell states assuming constant rates, then embedded them in a projection.

The cell type-specific RNA velocities (**Supplementary Figs. 9-12**) depict a highly directional landscape. The corresponding protein velocities (**Supplementary Figs. 13-16**) are much noisier as a result of sparser data collection (dozens of proteins vs. thousands of genes). We used a Gaussian kernel to determine the net velocities at regular grid points. The RNA and protein velocity fields (**Supplementary Figs. 17-20**), suggest that alignment between the two is strongly associated with cell type. The combination of RNA and protein velocities reveals the curvature of the cell state landscape. With respect to reference frame of unspliced transcripts, the protein inferences correspond to second-order protein *acceleration* of unspliced transcript counts. We visualize cell movement in the spliced mRNA using a Bezier curve calculated from three points corresponding to past, present, and future.

The protein acceleration landscape for the CITE-seq data shows a diversity of dynamics (**Fig. 2a**). B cells and a subset of T lymphocytes exhibit strong protein acceleration. This behavior may reflect recent findings^12^ that describe mRNA transcript “pile-up” due to heavily suppressed translation in naïve CD4+ T cells. The unidirectional monocyte velocity suggests response or plasticity^13^. However, due to the imperfect separation of cell types in the embedding, we caution against over-interpretation of aggregated velocities.

**Figure 2.**
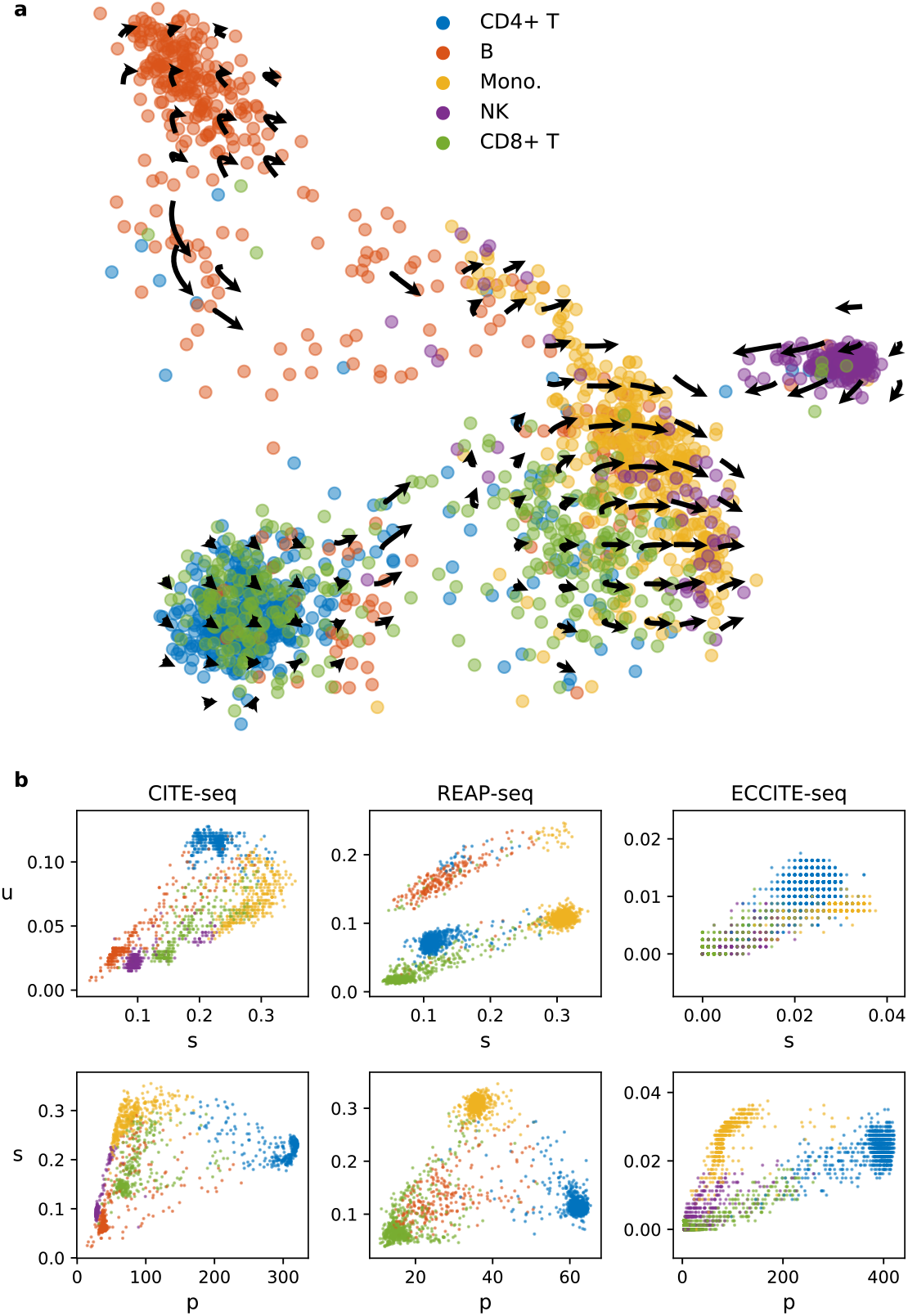
Protein acceleration visualization. (a) CITE-seq PBMC protein acceleration, visualized on a grid in principal component space. (b) Splicing and translation phase portraits of CD4 in three of the analyzed PBMC datasets.

REAP-seq results show similar T lymphocyte circulation and partitioning into static and mobile populations (**Supplementary Fig. 21**), albeit with noisier data than CITE-seq. ECCITE-seq yields sparse velocity landscapes (**Supplementary Figs. 22-23**), which result from the very shallow sequencing of spliced transcripts: we confirmed that ECCITE-seq captures 1-2 orders of magnitude fewer RNA molecules per cell than CITE-seq or REAP-seq, which is consistent with Fig. 1b of Mimitou et al.^5^ (**Supplementary Fig. 24**). In addition to using genes with linear behavior to infer velocity, we qualitatively confirmed consistency between datasets for the gene CD4, which does not follow linear behavior (**Fig. 2b**).

Our qualitative protein acceleration framework does not attempt to account for regulatory differences between cell types. Future work may involve better measurements that enable inference of parameters for a more gene-specific model of regulation that takes into account specific biochemical modulators. Current protein quantification protocols are adapted for histological markers on the cell surface; technology that can quantify dissolved protein could aid in more extensive studies of cell state and kinetics.

## Acknowledgements

G.G., V.S., and L.P. were partially funded by NIH U19MH114830. We thank the authors of Mimitou et al. 2019 for providing *velocyto* pipeline outputs for ECCITE-seq datasets.

## Author contributions

G.G. implemented the method and performed analysis under the supervision of V.S. V.S. and L.P. developed the protein acceleration extension of RNA velocity. G.G., V.S., and L.P. interpreted results and wrote the manuscript.

## Methods

Data were acquired from the Gene Expression Omnibus (GEO). We used the following matched RNA/protein datasets. CITE-seq: GSM2695381, GSM2695382. REAP-seq: GSM2685238, GSM2685243. ECCITE-seq ctrl: GSM3596095, GSM3596096. ECCITE-seq CTCL: GSM3596100, GSM3596101. Reads were aligned to the GRCh38 genome reference using the *Cell Ranger* pipeline with default settings whenever aligned sequence files were not already available through GEO. *Cell Ranger* 2.2.0 was used to align CITE-seq reads; *Cell Ranger* 3.0.0 was used for ECCITE-seq.

The aligned data were processed with the velocyto command-line interface. The resulting velocyto loom files with spliced and unspliced RNA counts assigned to each gene-cell pair were compared to the protein counts to identify common cells (*n* = 1780 cells for CITE-seq, 3158 for REAP-seq, 5084 for ECCITE-seq control, 5329 for ECCITE-seq CTCL). Protein counts were normalized to the median total protein number in each dataset. *k*-nearest neighbor imputation was performed on the logarithm of normalized protein counts to compute a graph of *k* nearest neighbors (*k* = 400 for CITE-seq, 800 for REAP-seq and ECCITE-seq) using the *scikit-learn* 0.20.0 Python package. For each cell, unspliced RNA, spliced RNA, and protein counts were assigned the mean value of the *k* neighbor cells. For ease of visualization, Louvain clustering was performed on the graph to identify cell clusters using the *louvain* 0.6.1 Python package. The ModularityVertexPartition model was used for CITE-seq and ECCITE-seq CTCL; RBERVertexPartition was used for REAP-seq and ECCITE-seq control. Cell types were manually assigned based on markers (**Supplementary Table 2**) reported in CITE-seq and REAP-seq publications^3,4^ (**Supplementary Figs. 25-28**). To calculate RNA velocity, the *velocyto* 0.17 workflow with default settings was used to fit extreme quantiles of the phase plot and extrapolate deviations from the equilibrium line. Normalized spliced expression was projected into an embedding for visualization. CITE-seq and REAP-seq data were embedded into the principal component space (PC2/PC3). ECCITE-seq data were embedded into a t-Distributed Stochastic Neighbor Embedding (t-SNE) calculated from the first 25 principal components. The *scikit-learn* package was used for PCA and t-SNE calculations. Transition probabilities to each cell’s embedding neighborhood of *m* cells were calculated, then aggregated for projection and visualization (*m* = 500 for CITE-seq, 200 for REAP-seq, 300 for ECCITE-seq). The embedding-specific future cell state was estimated by adding the projected displacement to the present cell state. Grid arrows were generated by applying a Gaussian kernel to the cell-specific velocities of *j* nearest neighbor cells (*j* = 100 for CITE-seq and REAP-seq, 500 for ECCITE-seq). The aggregated future state corresponding to each grid point was calculated by adding this RNA forward velocity vector to the grid point.

To calculate protein velocity, pairs of proteins and source spliced RNA consistent with the linear model hypothesis were manually selected to stand in for spliced and unspliced RNA. These were identically projected onto the same embedding (*m* = 499). Forward grid arrows were generated by applying a Gaussian kernel to the cell-specific velocities of *l* nearest neighbor cells (*l* = 200). The aggregated past state corresponding to each grid point was calculated by subtracting this protein forward velocity vector from the grid point. Curved arrows corresponding to the entire trajectory were produced by fitting a second-order Bézier curve to the sequence of past, present, and future states of each grid point using the *bezier* 0.9.0 Python package.

Scripts to reproduce the results of this paper are available at https://github.com/pachterlab/GSP_2019.

