## Supplementary Note, Tables, Figures. for "RNA velocity and protein acceleration from single-cell multiomics experiments"

### RNA velocity

We assume that the abundance of an RNA molecule is determined by the following differential equations:

$$\frac{du}{ds} = \alpha(t) - \beta_s(t) \cdot u;$$

$$\frac{ds}{dt} = \beta_s(t) \cdot u - \gamma_s(t) \cdot s.$$

The transcription, splicing, and degradation parameters $\alpha, \beta_s, \gamma_s$ are gene- and time-dependent. Neglecting the time dependence produces a simplified system for each gene:

$$\frac{du}{ds} = \alpha - \beta_s \cdot u;$$

$$\frac{ds}{dt} = \beta_s \cdot u - \gamma_s \cdot s.$$

Under the assumptions underlying the RNA velocity framework[6], $\beta_s$ is a common, gene-independent splicing parameter. Dividing by $\beta_s$ yields:

$$\frac{1}{\beta_s} \cdot \frac{du}{ds} = \frac{\alpha}{\beta_s} - u = \hat{\alpha} - u;$$

$$\frac{1}{\beta_s} \cdot \frac{ds}{dt} = u - \frac{\gamma_s \cdot s}{\beta_s} = u - \widehat{\gamma_s} \cdot s.$$

The equilibrium line is defined by $\frac{ds}{dt} = 0 = u - \widehat{\gamma_s} \cdot s$, yielding $\widehat{\gamma_s} = u/s$. Therefore, the rate of change in spliced counts scales with deviation from equilibrium. We estimate the normalized degradation parameter $\widehat{\gamma_s}$ using imputed raw (non-normalized) U and S counts, and apply extreme quantile fitting[6]. Extrapolation is a first-order forward estimate, calculated using a constant velocity.

### Protein velocity

Protein production and degradation are governed by the following equation:

$$\frac{dp}{dt} = \beta_p(t) \cdot s - \gamma_p(t) \cdot p.$$

Assuming constant parameters, this results in:

$$\frac{dp}{dt} = \beta_p \cdot s - \gamma_p \cdot p;$$

$$\frac{1}{\beta_p} \cdot \frac{dp}{dt} = s - \frac{\gamma_p \cdot p}{\beta_p} = s - \widehat{\gamma_p} \cdot p.$$

Following the RNA velocity framework, we can use a similar approximation and assume the translation rate is gene-independent. The corresponding equilibrium line is defined by $\frac{dp}{dt} = 0 = s - \widehat{\gamma_p} \cdot p$, yielding $\widehat{\gamma_p} = s/p$. The rate of change in protein counts scales with deviation from this equilibrium. We estimate the normalized degradation parameter $\widehat{\gamma_p}$ using imputed raw (non-normalized) S and P counts, and apply extreme quantile fitting. As with RNA velocity, extrapolation is first-order; however, protein velocity is a backward estimate.

### Quantitative considerations

Since both splicing[14] and translation[15] are very tightly regulated in eukaryotes, the assumption of constant rates across all cells and genes means the extrapolation step is qualitative. Furthermore, the sparsity of the protein data amplifies deviations.

The two resulting differential equations are normalized to different parameters, but we do not weigh the extrapolation results differently when constructing the past and future estimates. This approach is consistent with the RNA velocity framework, whose extrapolated state is a mean neighbor cell transition target; scaling by velocity magnitude is optional.

Further, for qualitative purposes, the timescales of translation and splicing are comparable. From mean estimates for human genes, we take the translation initiation time to be 7 s and the elongation time per codon to be 87 ms[16]. For the seven proteins used for protein velocity estimation in the CITE-seq dataset, the expected average translation timescale is 0.5 – 1 min. Sources are equivocal on the rate of splicing; recent estimates yield 0.5 – 5 min[17], comparable to translation.

### Supplementary Tables

| CITE-seq | REAP-seq | ECCITE-seq |
|---|---|---|
| CD3D | ITGAM | ITGAX |
| CD8A | HLA-DRA | CD2 |
| CD2 | CD8A | DPP4 |
| FCGR3A | CD8B | CD27 |
| CD14 | IL7R | CD28 |
| ITGAX | PTPRC | CD3E |
| CD19 | CD27 | CD5 |
| | CD9 | SELL |
| | CD19 | CD7 |
| | CD40 | CD8A |
| | CD4D | IL7R |
| | MS4A1 | HLA-DRA |
| | CD14 | |

**Supplementary Table 1**. Genes used for protein velocity estimation.

| CD4+ T | B | Monocytes | NK | CD8+ T |
|---|---|---|---|---|
| CD4 | CD45RA | CD11b | CD56 | CD8 |
| CD3 | CD19 | CD11c | | |
| | HLA-DR | CD14 | | |
| | | HLA-DR | | |

**Supplementary Table 2.** Canonical cell surface markers used for cell type identification[3,4].

### Supplementary Figures

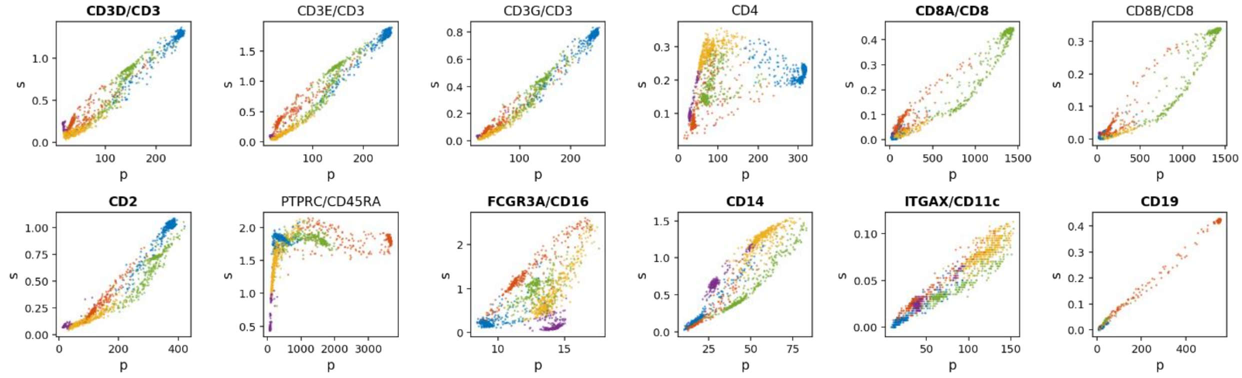


**Supplementary Figure 1.** CITE-seq imputed RNA/protein phase plots. Pairs used for protein velocity estimation are shown in bold. Color identifies cell type (blue: CD4+ T, red: B, yellow: monocytes, green: CD8+ T, purple: natural killer).

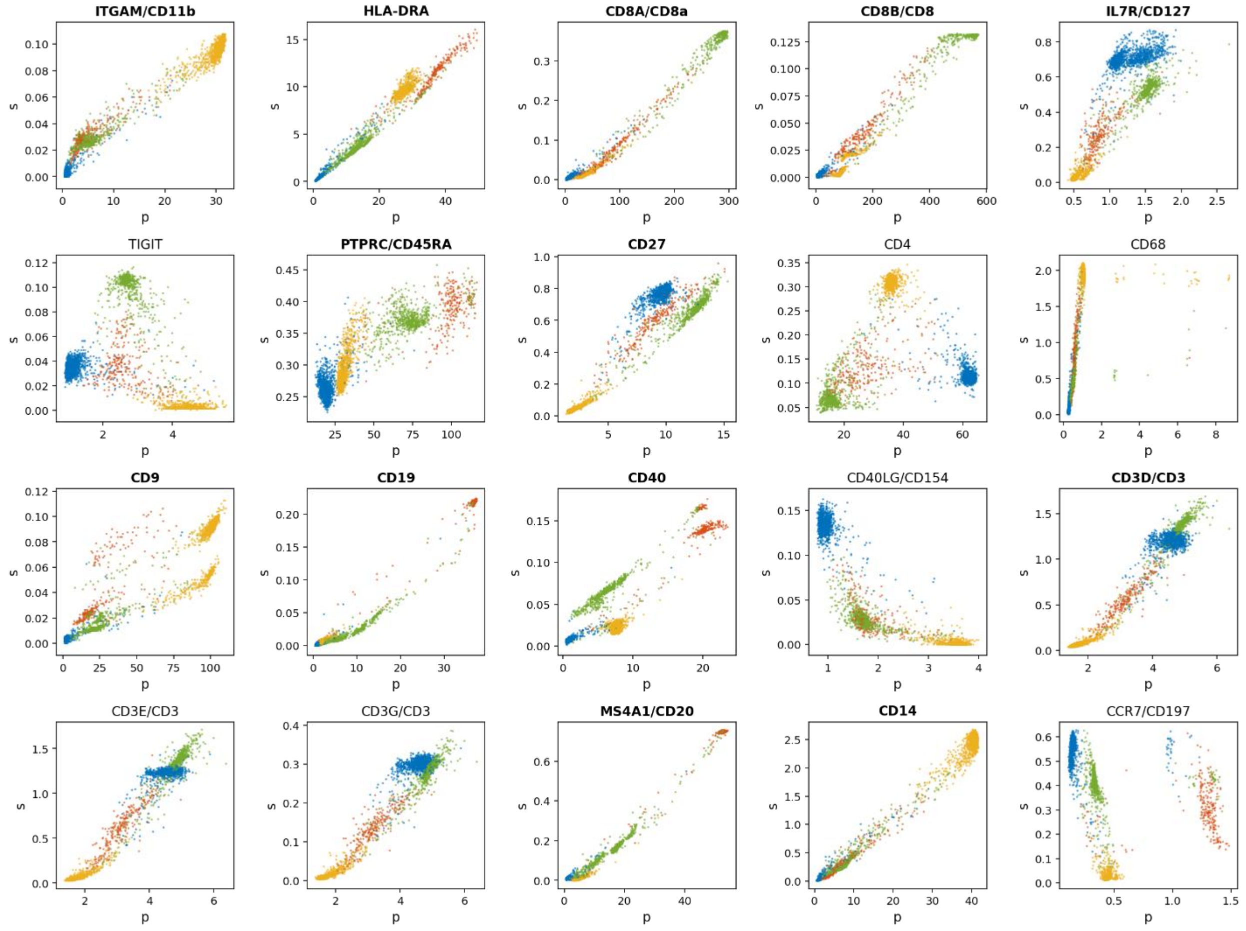


**Supplementary Figure 2.** REAP-seq imputed RNA/protein phase plots. Pairs used for protein velocity estimation shown in bold. Color identifies cell type (blue: CD4+ T, red: B, yellow: monocytes, green: CD8+ T).

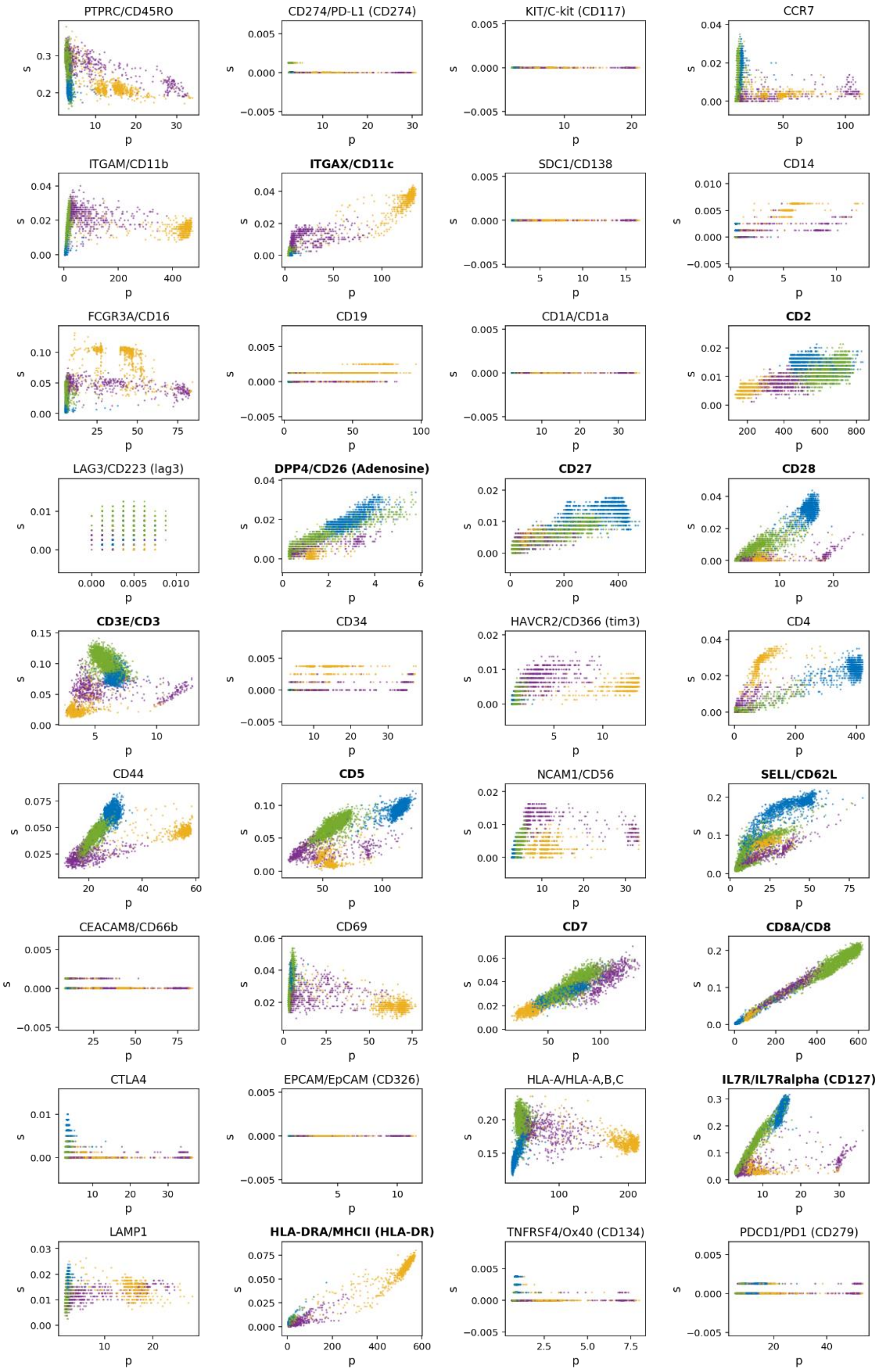


**Supplementary Figure 3.** ECCITE-seq ctrl imputed RNA/protein phase plots. Pairs used for protein velocity estimation shown in bold. Color identifies cell type (blue: CD4+ T, yellow: monocytes, green: CD8+ T, purple: natural killer).

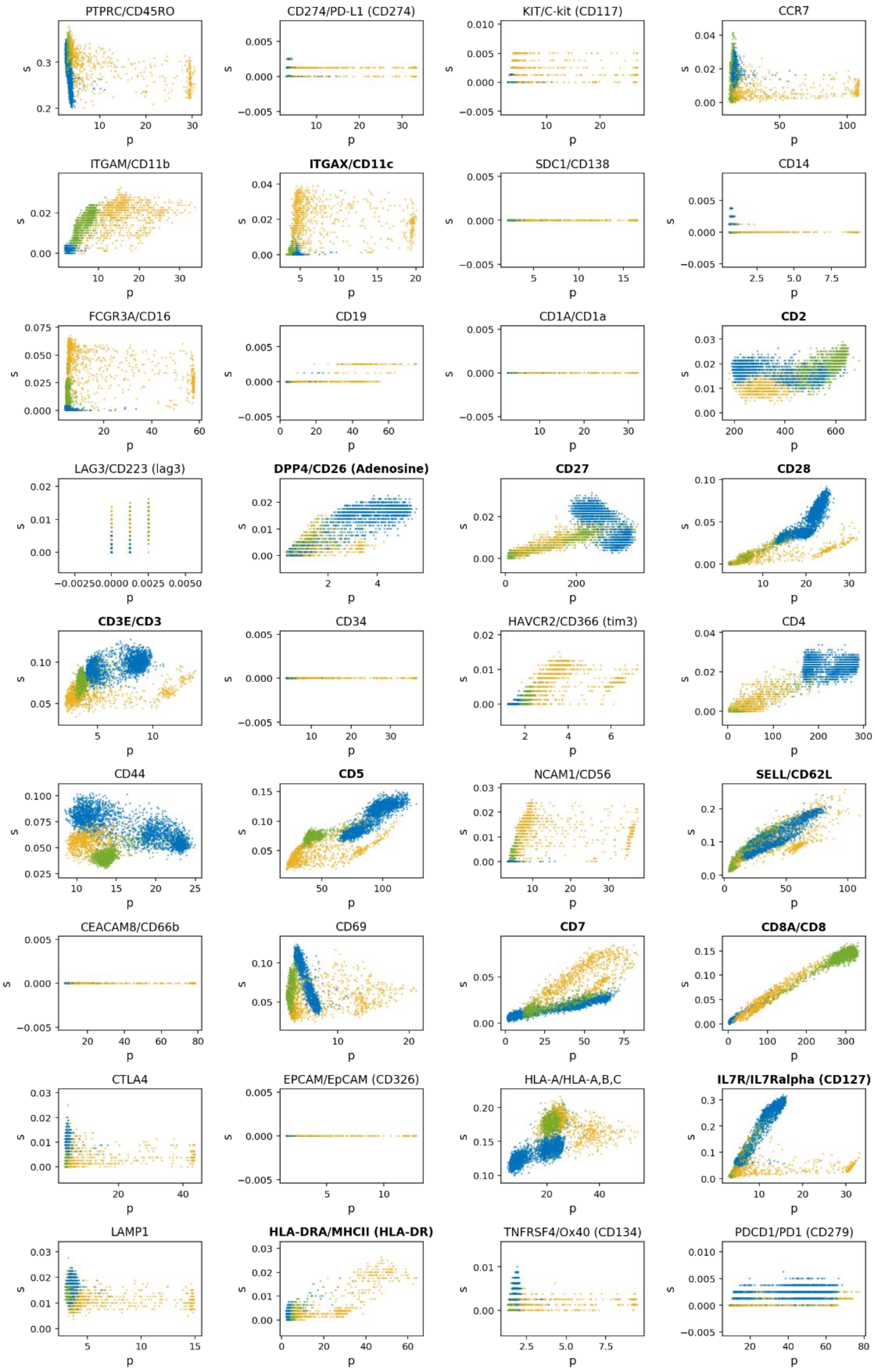


**Supplementary Figure 4.** ECCITE-seq CTCL imputed RNA/protein phase plots. Pairs used for protein velocity estimation shown in bold. Color identifies cell type (blue: CD4+ T, yellow: monocytes, green: CD8+ T).

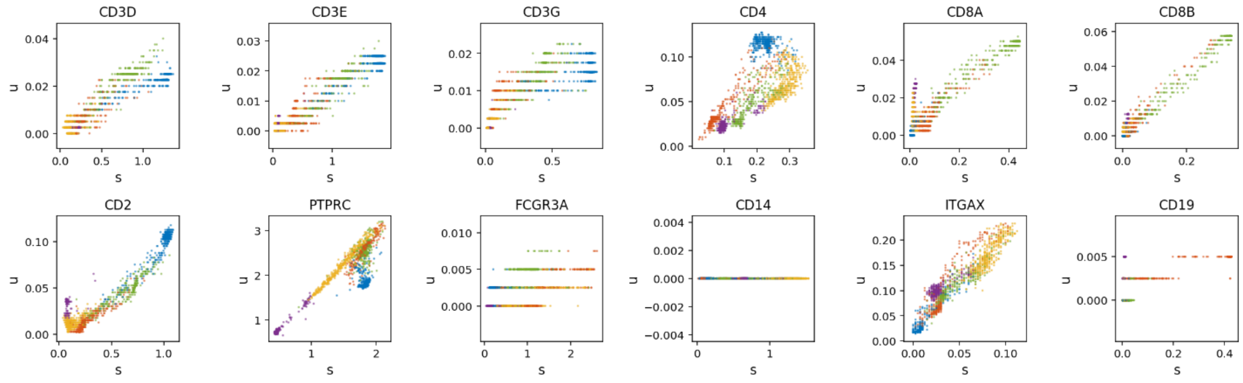


**Supplementary Figure 5.** CITE-seq imputed spliced/unspliced RNA phase plots. Only immunoglobulin-coding RNA are shown. Color identifies cell type (blue: CD4+ T, red: B, yellow: monocytes, green: CD8+ T, purple: natural killer).

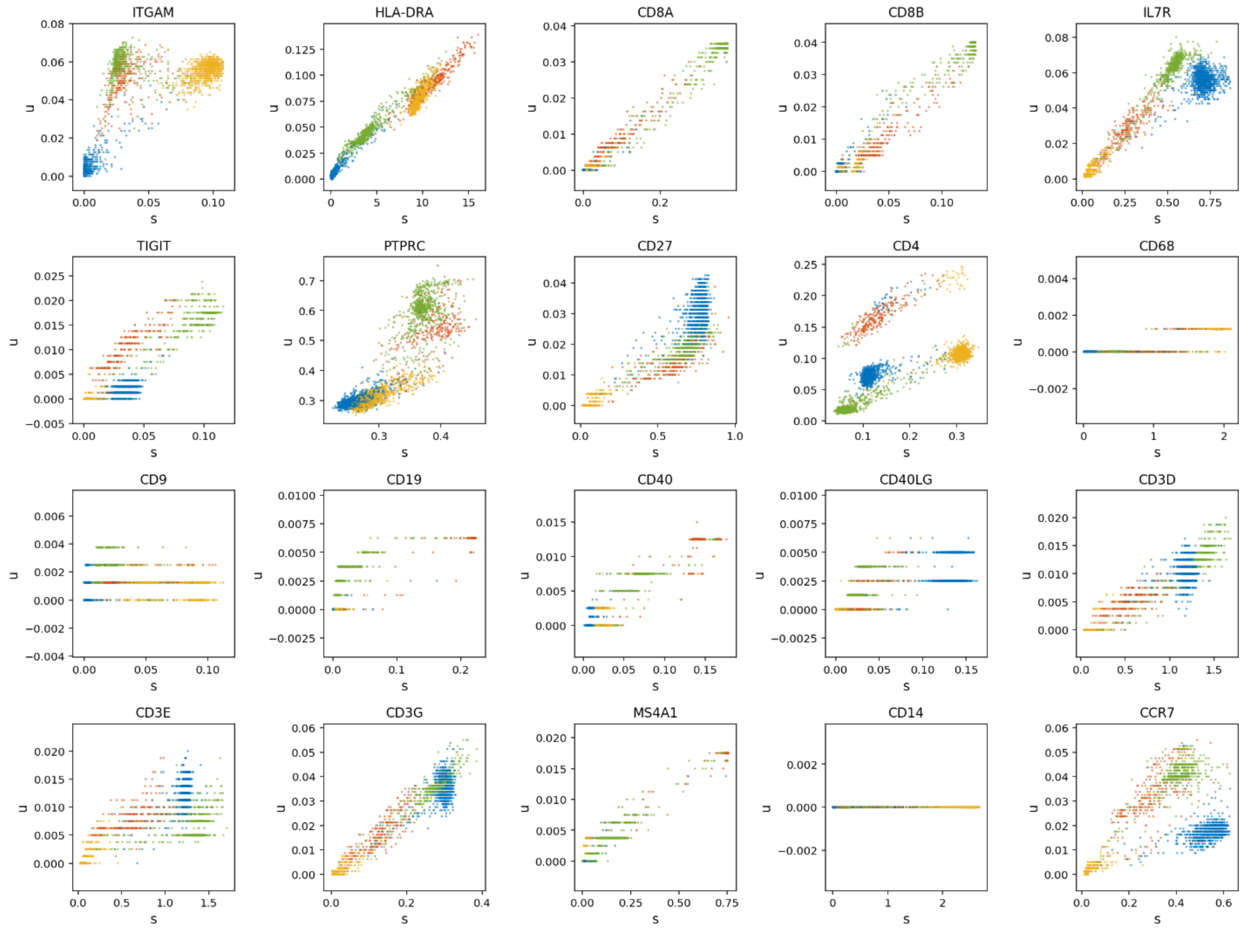


**Supplementary Figure 6.** REAP-seq imputed spliced/unspliced RNA phase plots. Only immunoglobulin-coding RNA are shown. Color identifies cell type (blue: CD4+ T, red: B, yellow: monocytes, green: CD8+ T).

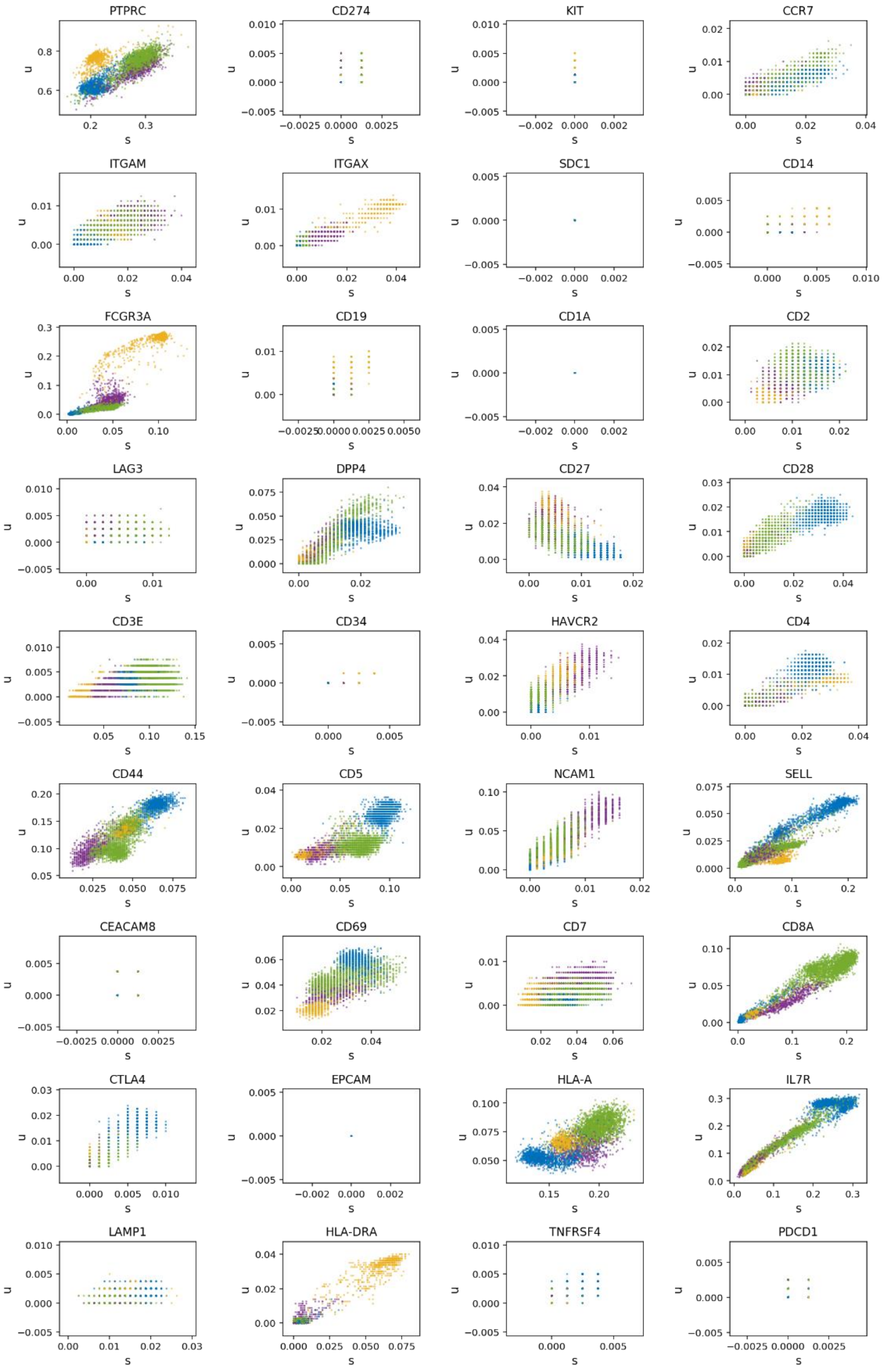


**Supplementary Figure 7.** ECCITE-seq control (ctrl) imputed spliced/unspliced RNA phase plots. Only immunoglobulin-coding RNA are shown. Color identifies cell type (blue: CD4+ T, yellow: monocytes, green: CD8+ T, purple: natural killer).

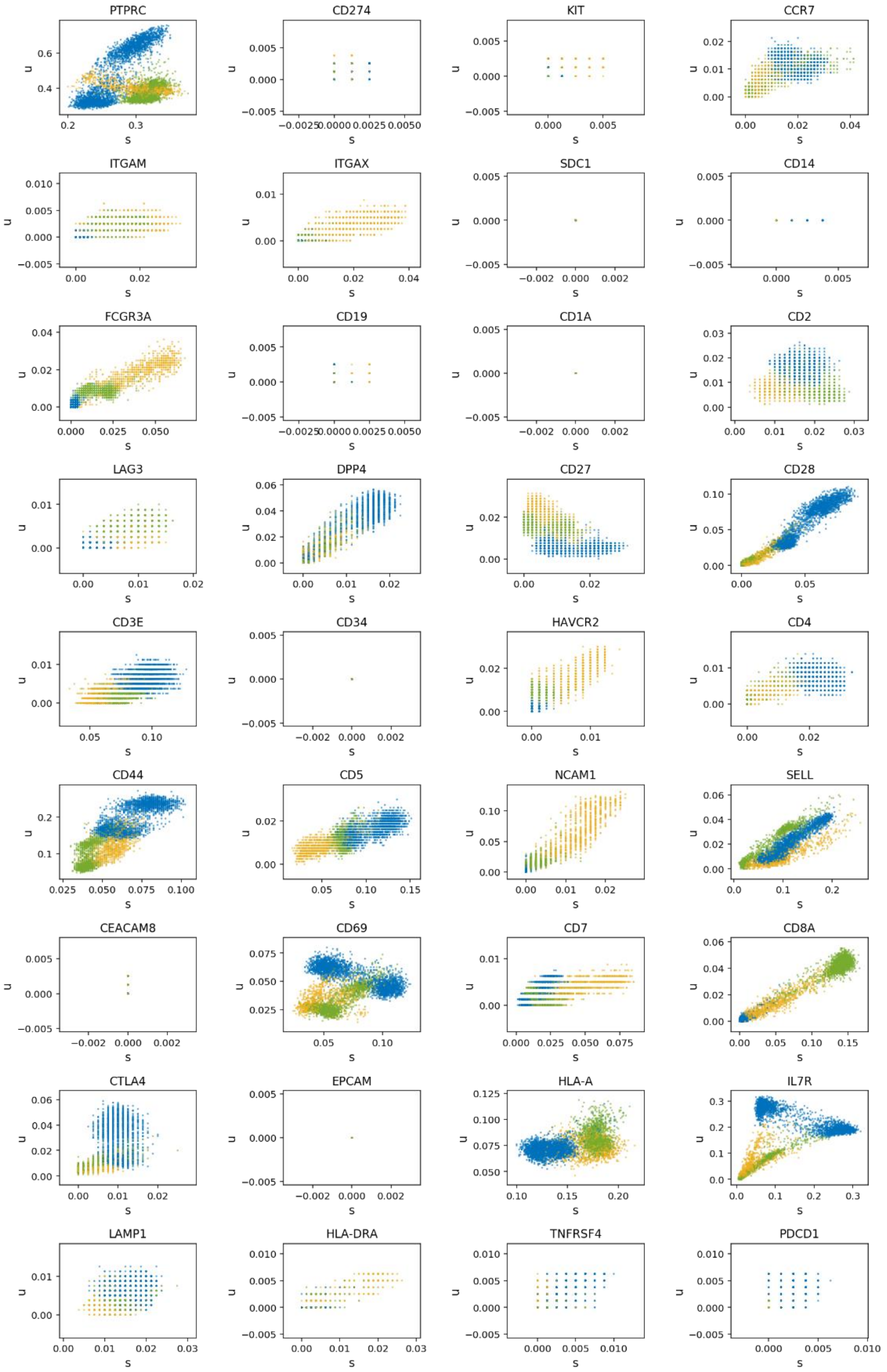


**Supplementary Figure 8.** ECCITE-seq CTCL patient (CTCL) imputed spliced/unspliced RNA phase plots. Only immunoglobulin-coding RNA are shown. Color identifies cell type (blue: CD4+ T, yellow: monocytes, green: CD8+ T).

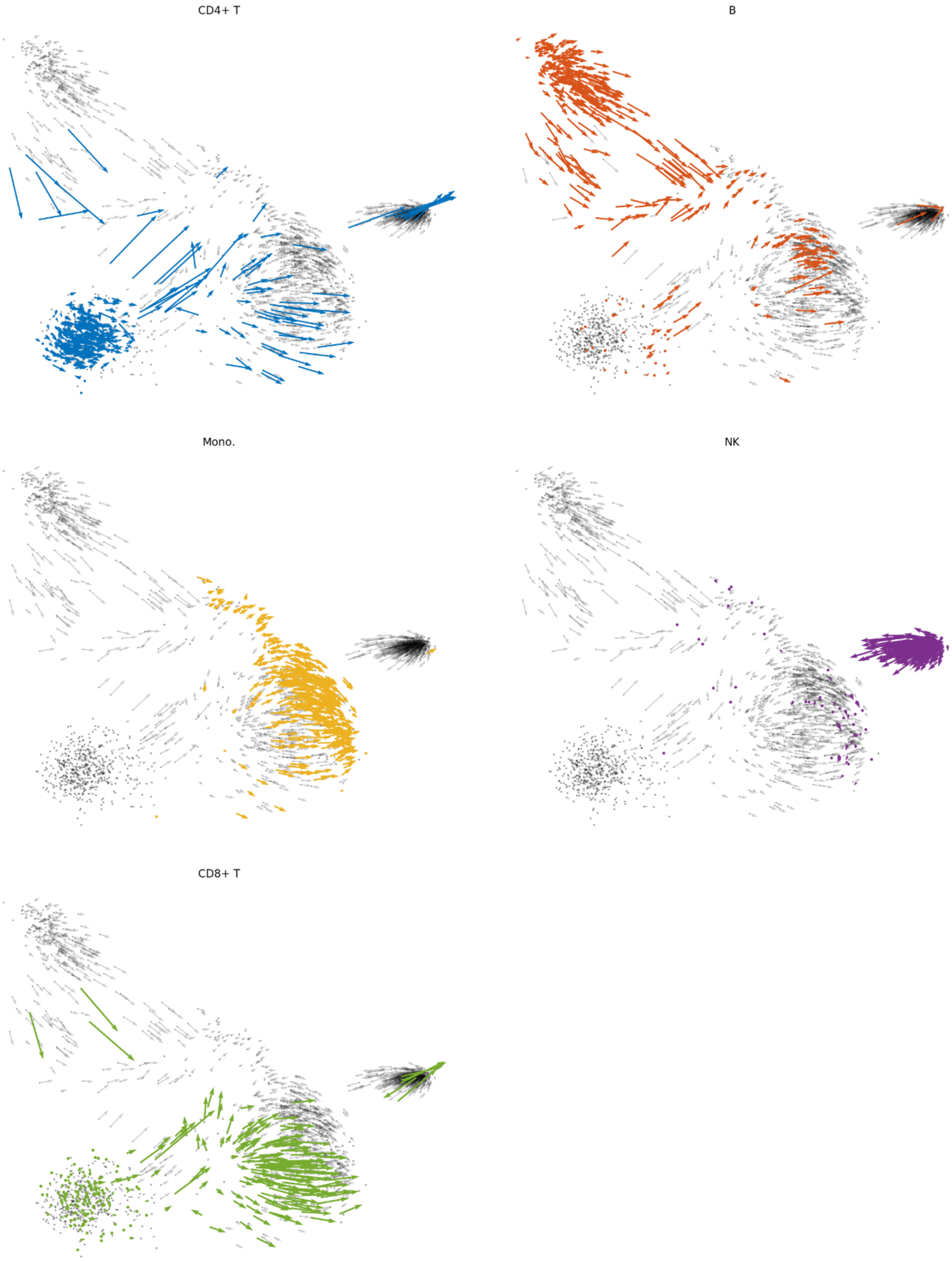


**Supplementary Figure 9**. CITE-seq cell RNA velocities, distinguished by cell type. Color identifies cell type (blue: CD4+ T, red: B, yellow: monocytes, green: CD8+ T, purple: natural killer).

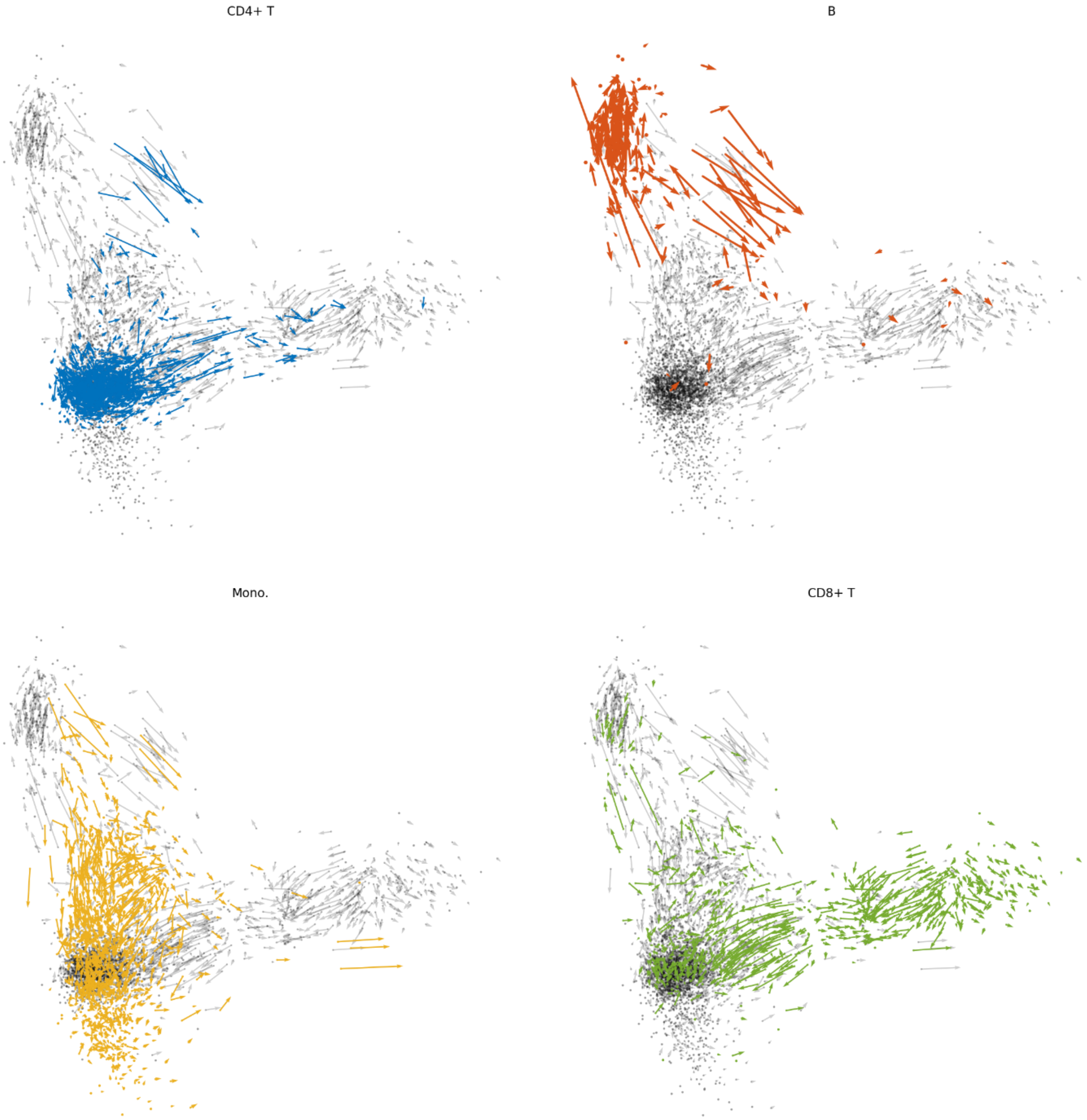


**Supplementary Figure 10**. REAP-seq cell RNA velocities, distinguished by cell type. Color identifies cell type (blue: CD4+ T, red: B, yellow: monocytes, green: CD8+ T).

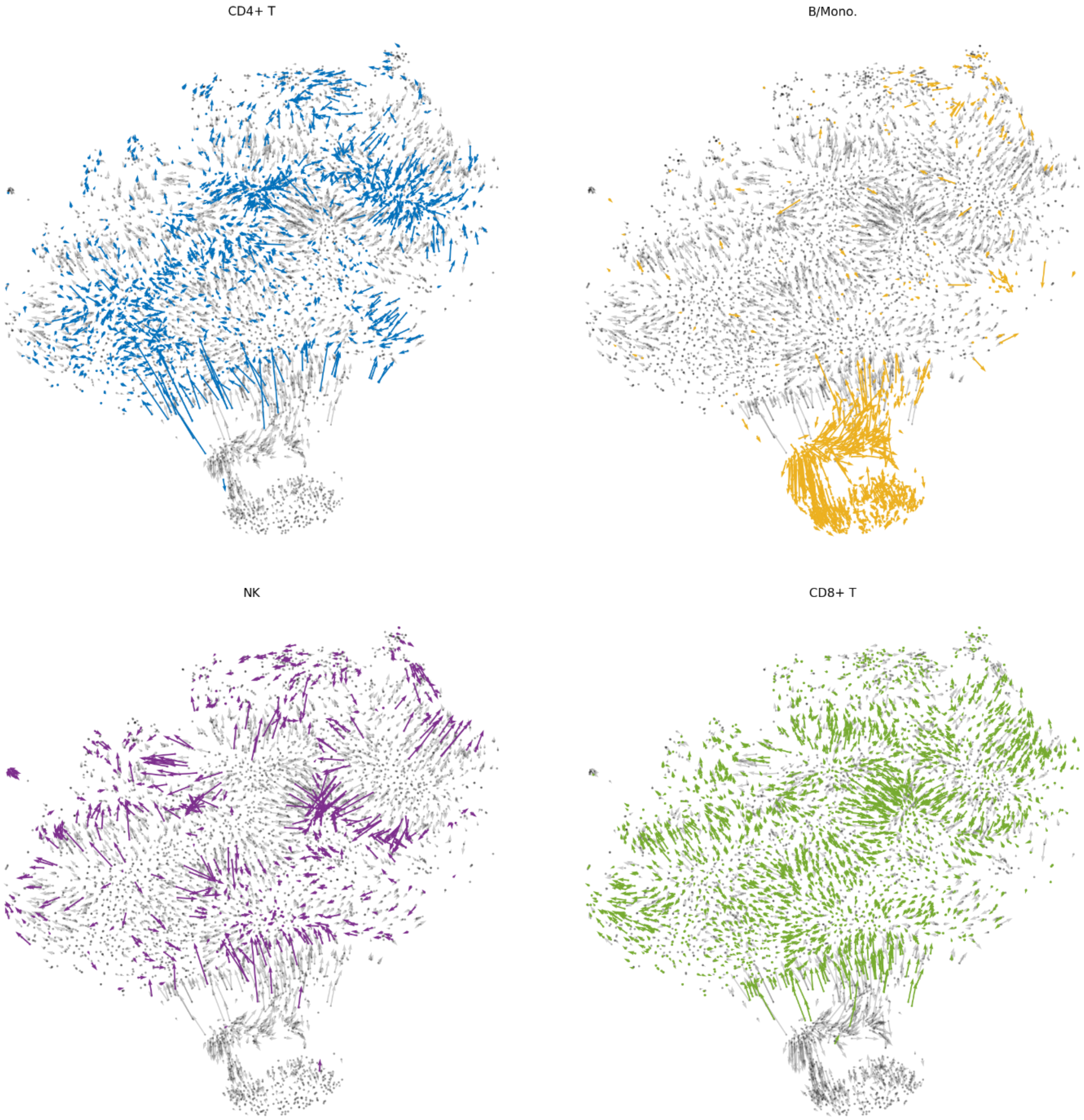


**Supplementary Figure 11.** ECCITE-seq ctrl cell RNA velocities, distinguished by cell type. Color identifies cell type (blue: CD4+ T, yellow: monocytes, green: CD8+ T, purple: natural killer).

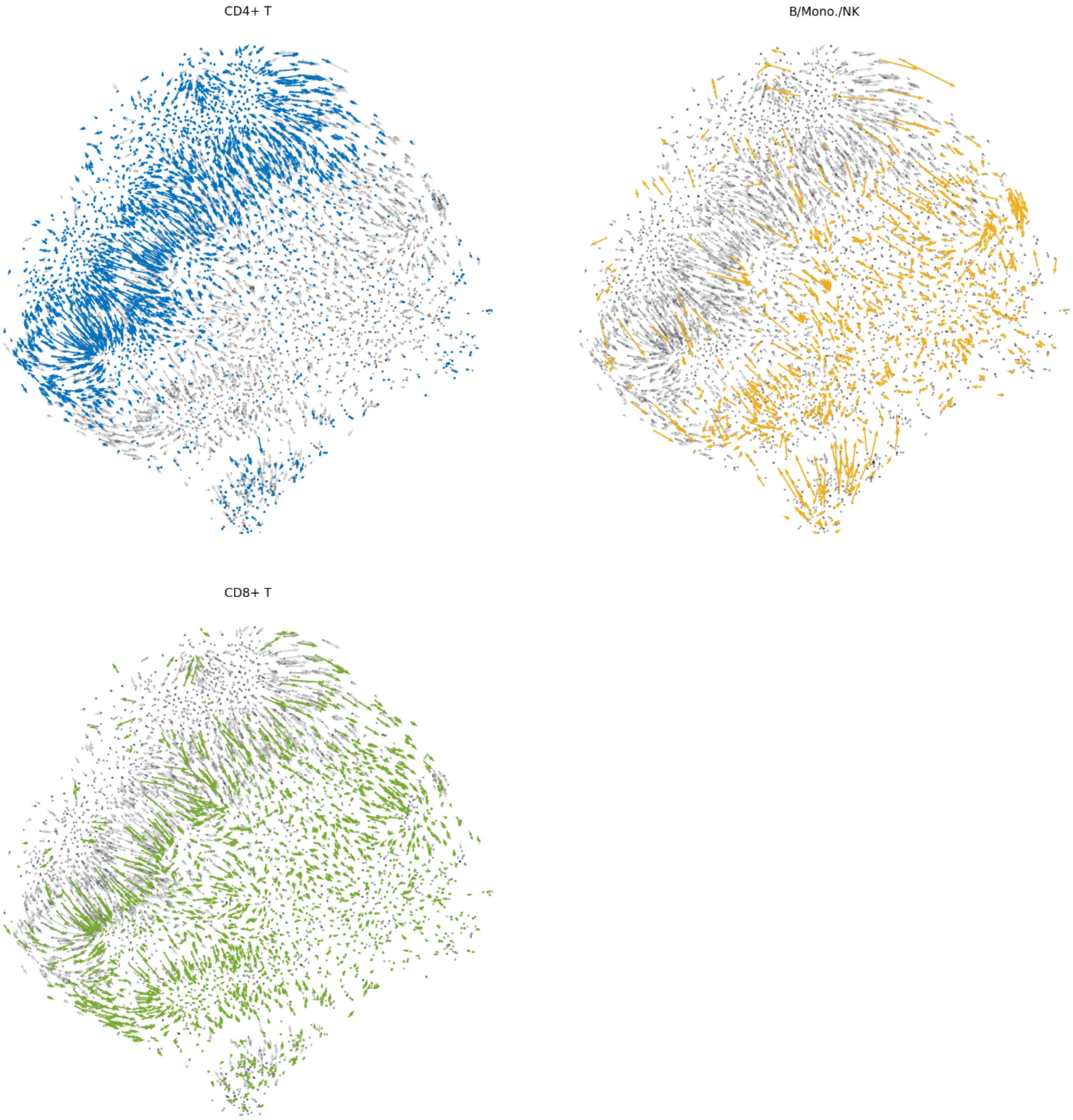


**Supplementary Figure 12.** ECCITE-seq CTCL cell RNA velocities, distinguished by cell type. Color identifies cell type (blue: CD4+ T, yellow: monocytes, green: CD8+ T).

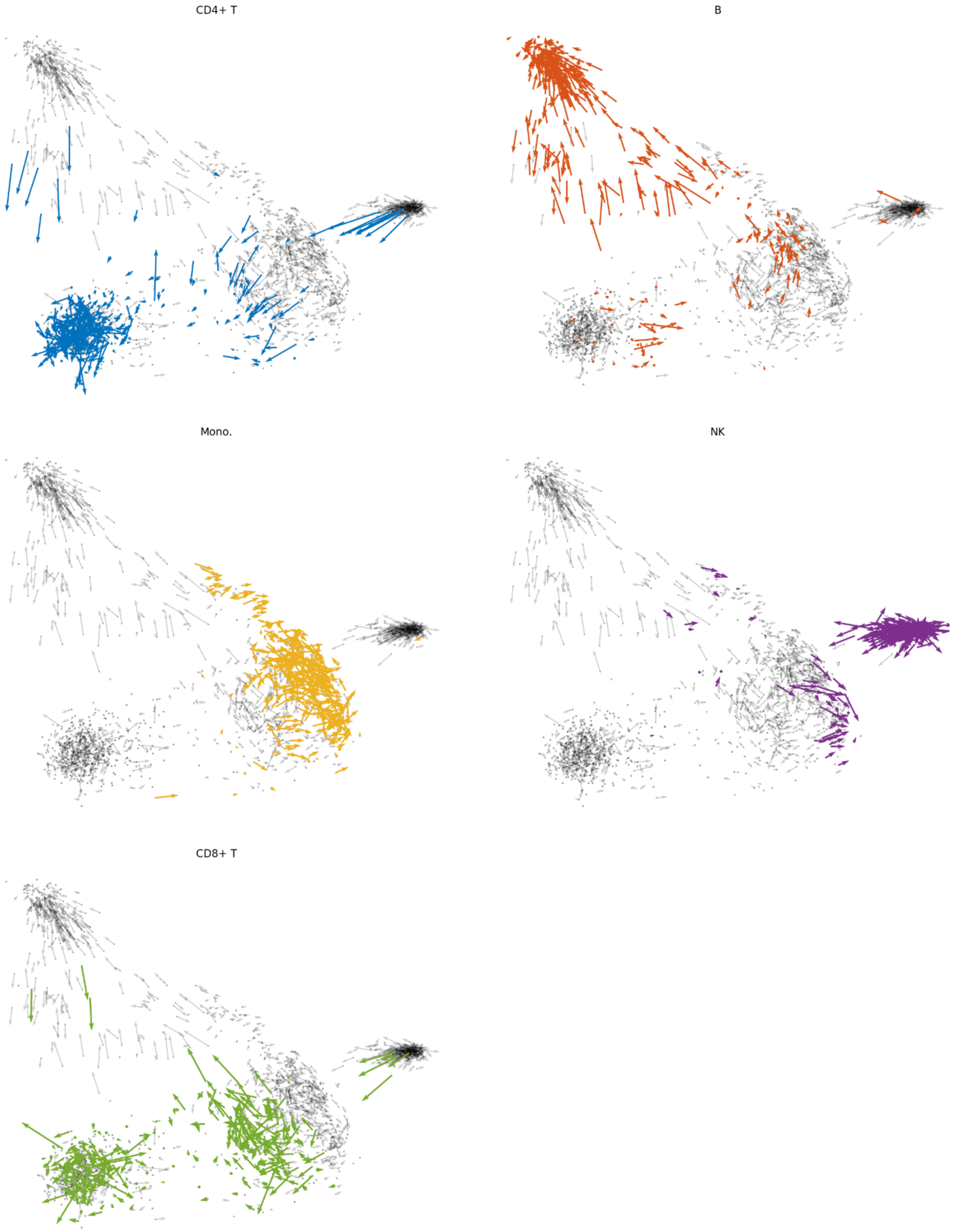


**Supplementary Figure 13**. CITE-seq cell protein velocities, distinguished by cell type. Color identifies cell type (blue: CD4+ T, red: B, yellow: monocytes, green: CD8+ T, purple: natural killer).

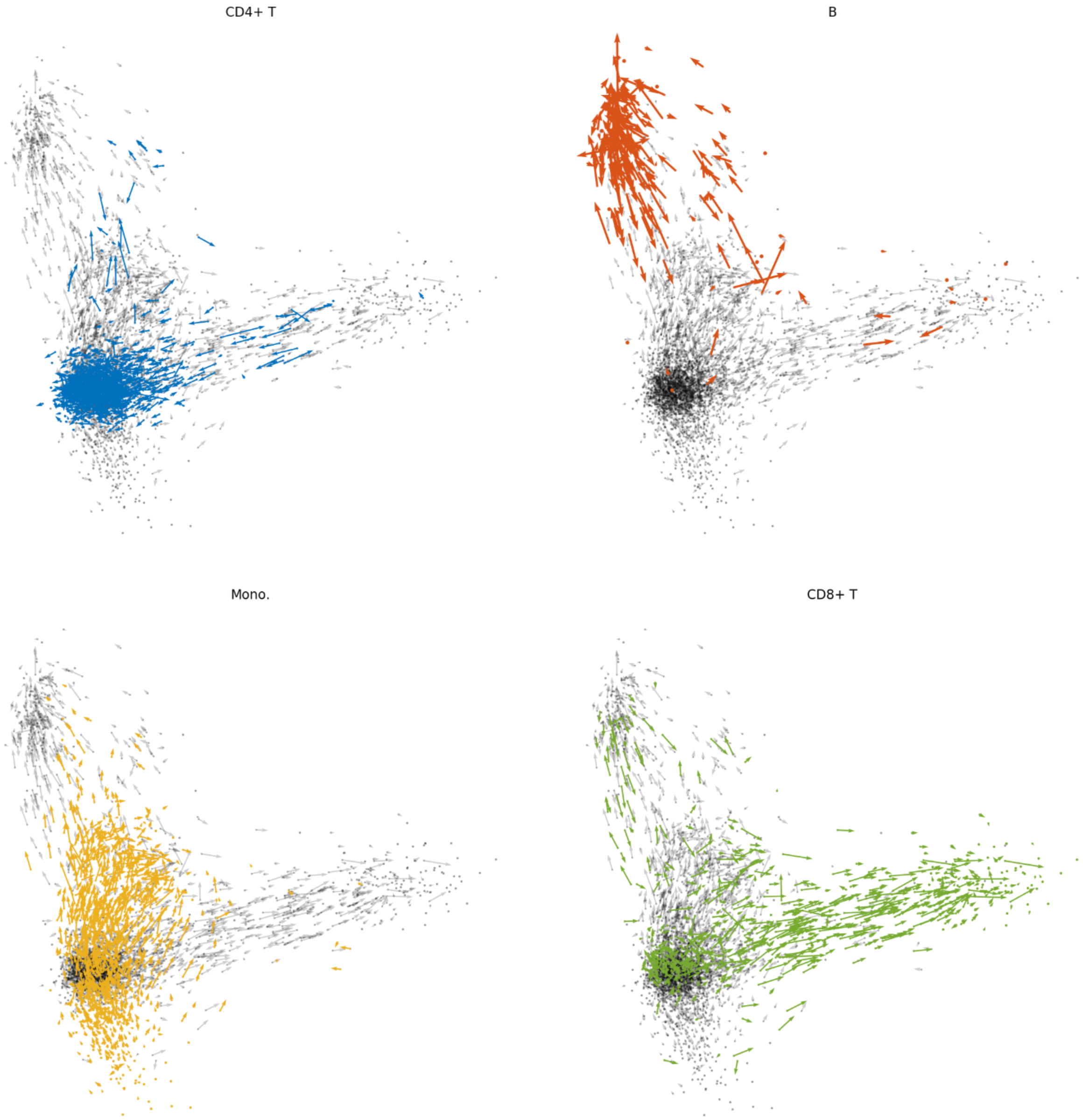


**Supplementary Figure 14**. REAP-seq cell protein velocities, distinguished by cell type. Color identifies cell type (blue: CD4+ T, red: B, yellow: monocytes, green: CD8+ T).

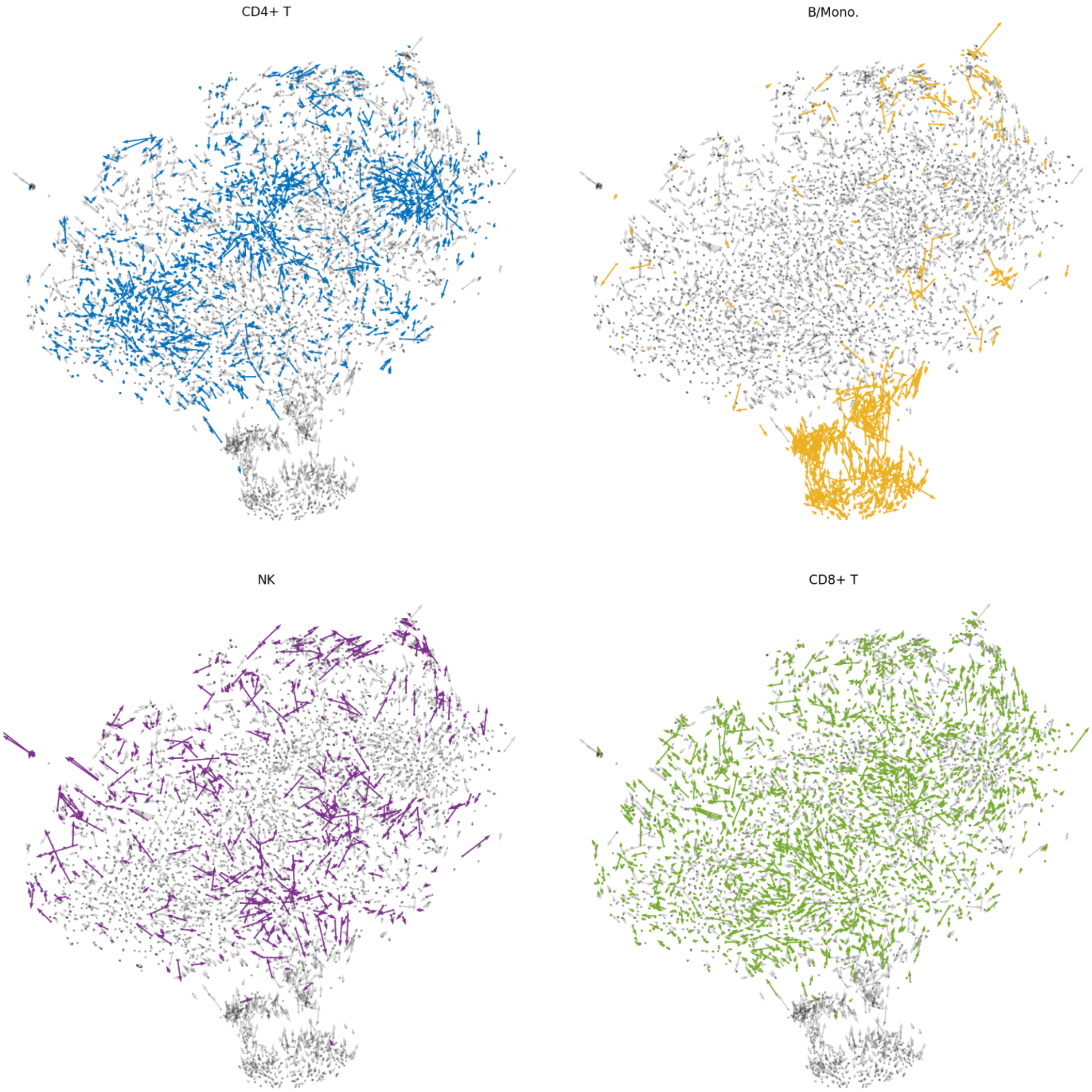


**Supplementary Figure 15.** ECCITE-seq ctrl cell protein velocities, distinguished by cell type. Color identifies cell type (blue: CD4+ T, yellow: monocytes, green: CD8+ T, purple: natural killer).

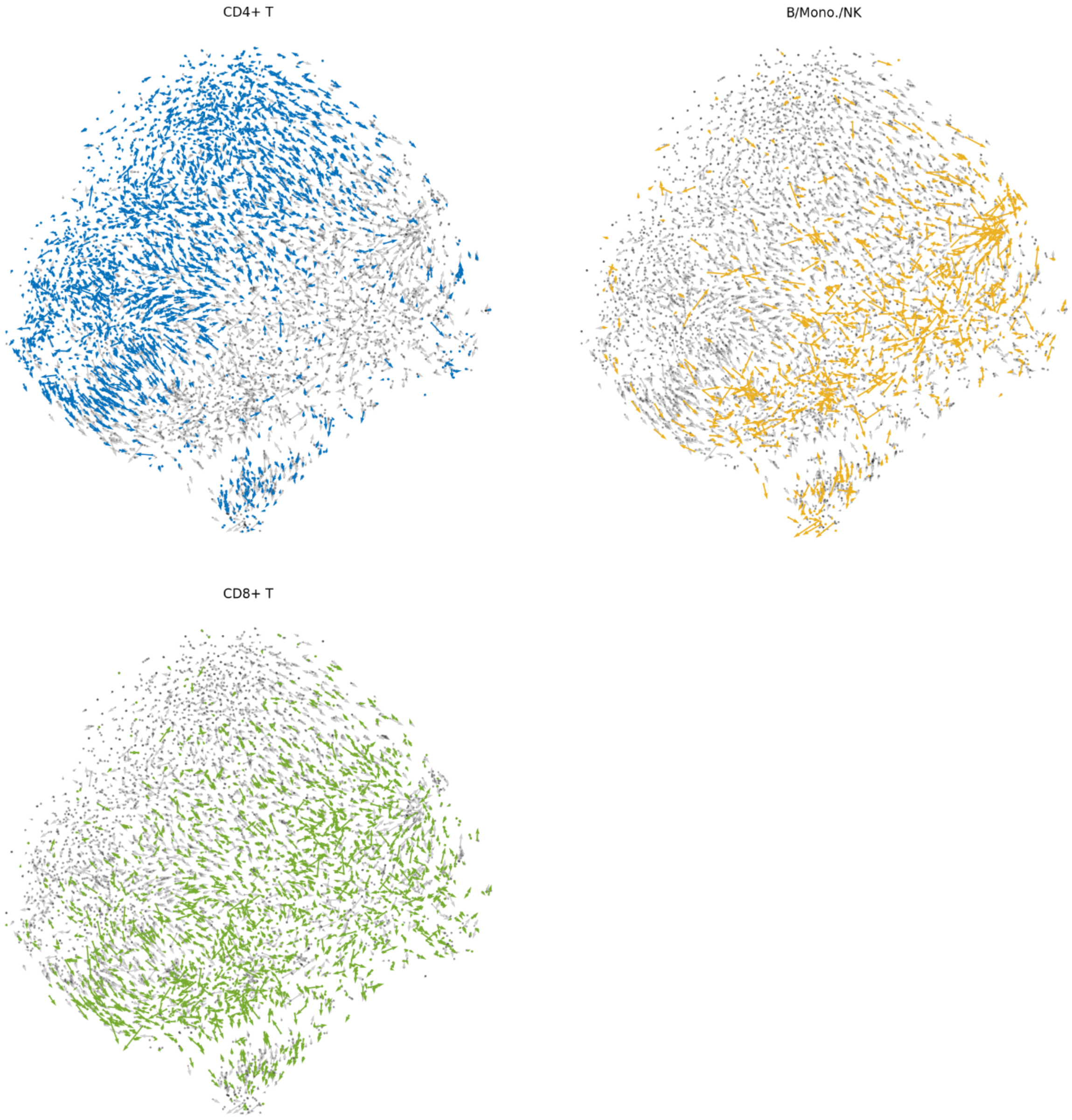


**Supplementary Figure 16.** ECCITE-seq CTCL cell protein velocities, distinguished by cell type. Color identifies cell type (blue: CD4+ T, yellow: monocytes, green: CD8+ T).

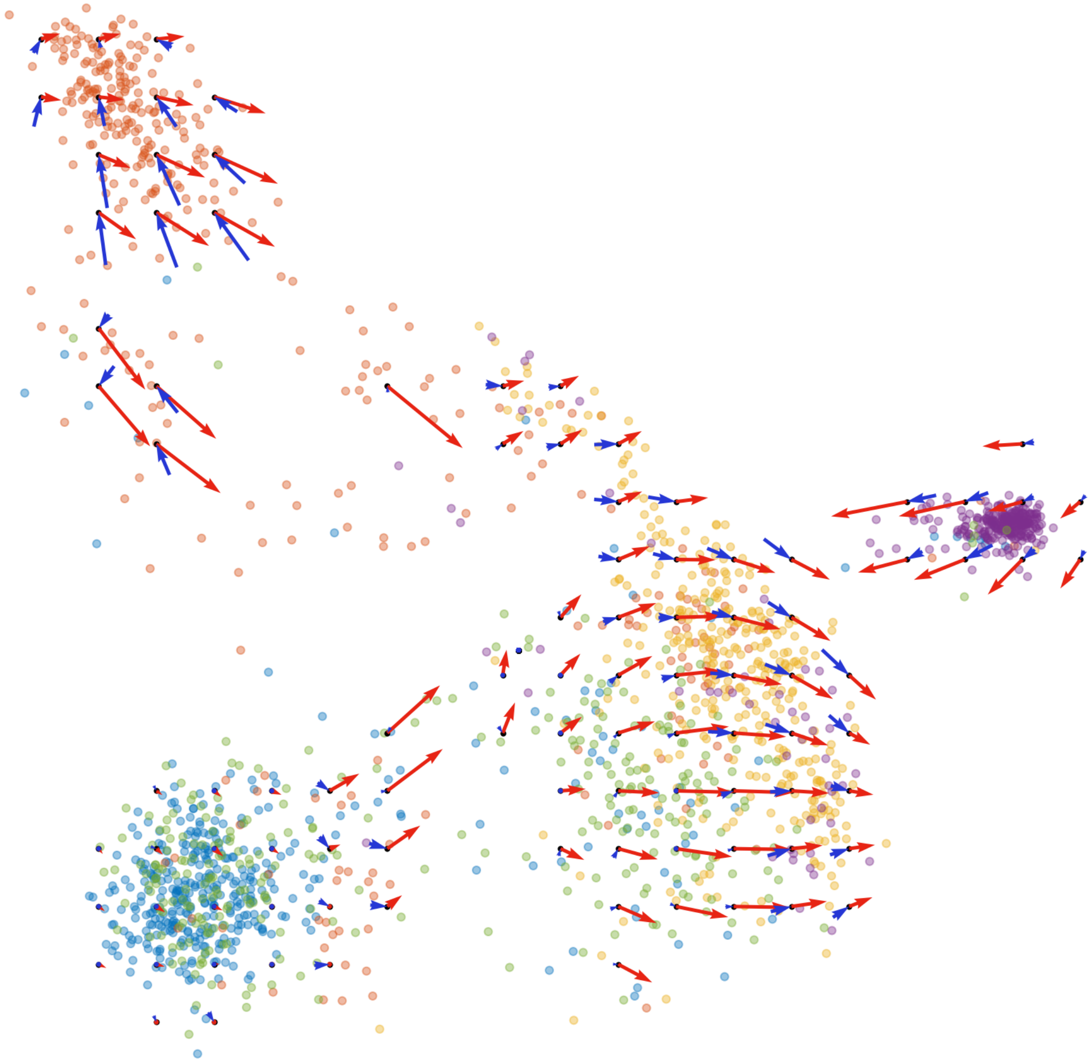

**Supplementary Figure 17**. CITE-seq velocity fields visualized on a grid. Arrow color identifies velocity estimate (RNA: red, protein: blue). Dot color identifies cell type (blue: CD4+ T, red: B, yellow: monocytes, green: CD8+ T, purple: natural killer).

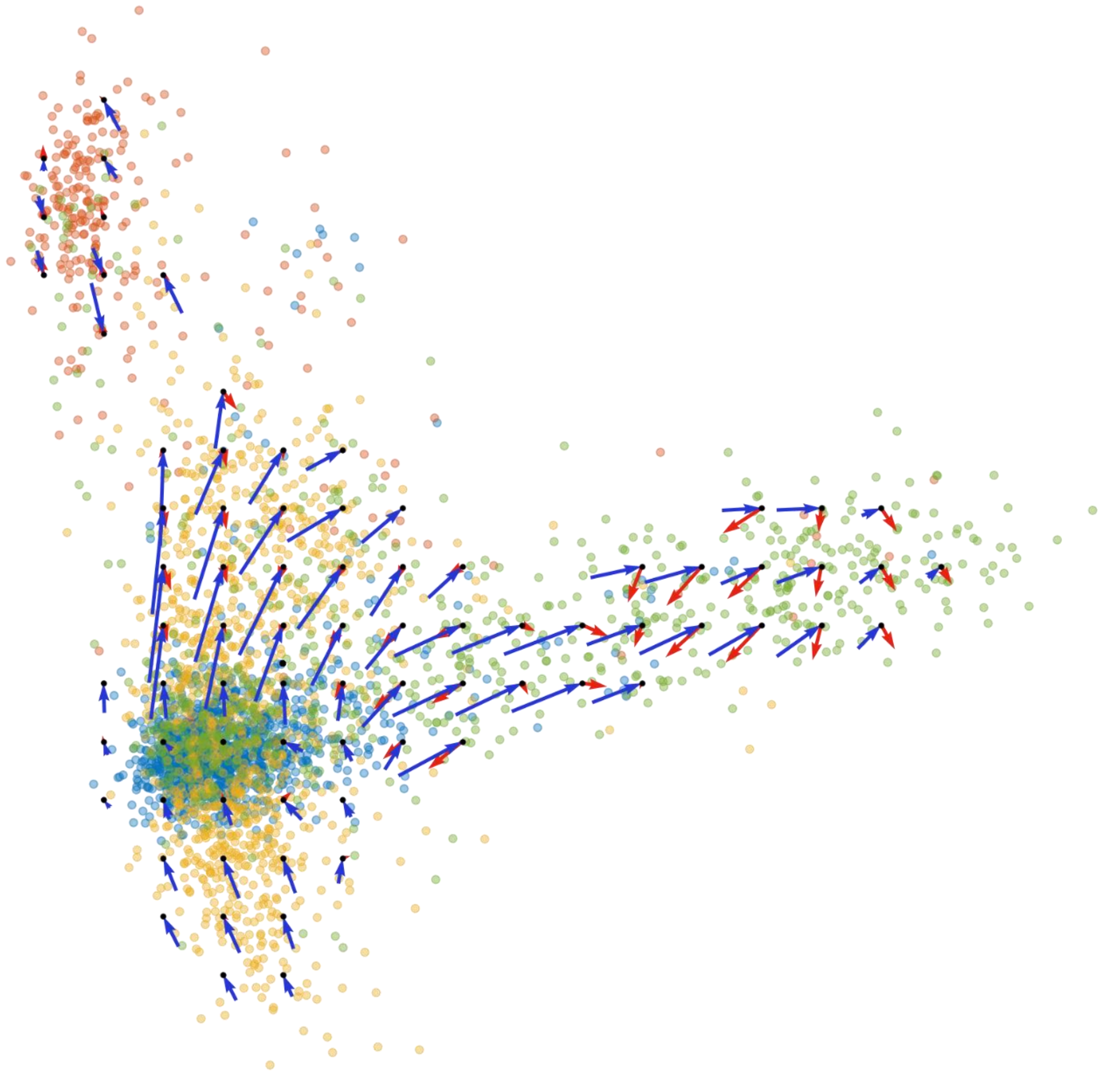

**Supplementary Figure 18.** REAP-seq velocity fields visualized on a grid. Arrow color identifies velocity estimate (RNA: red, protein: blue). Dot color identifies cell type (blue: CD4+ T, red: B, yellow: monocytes, green: CD8+ T).

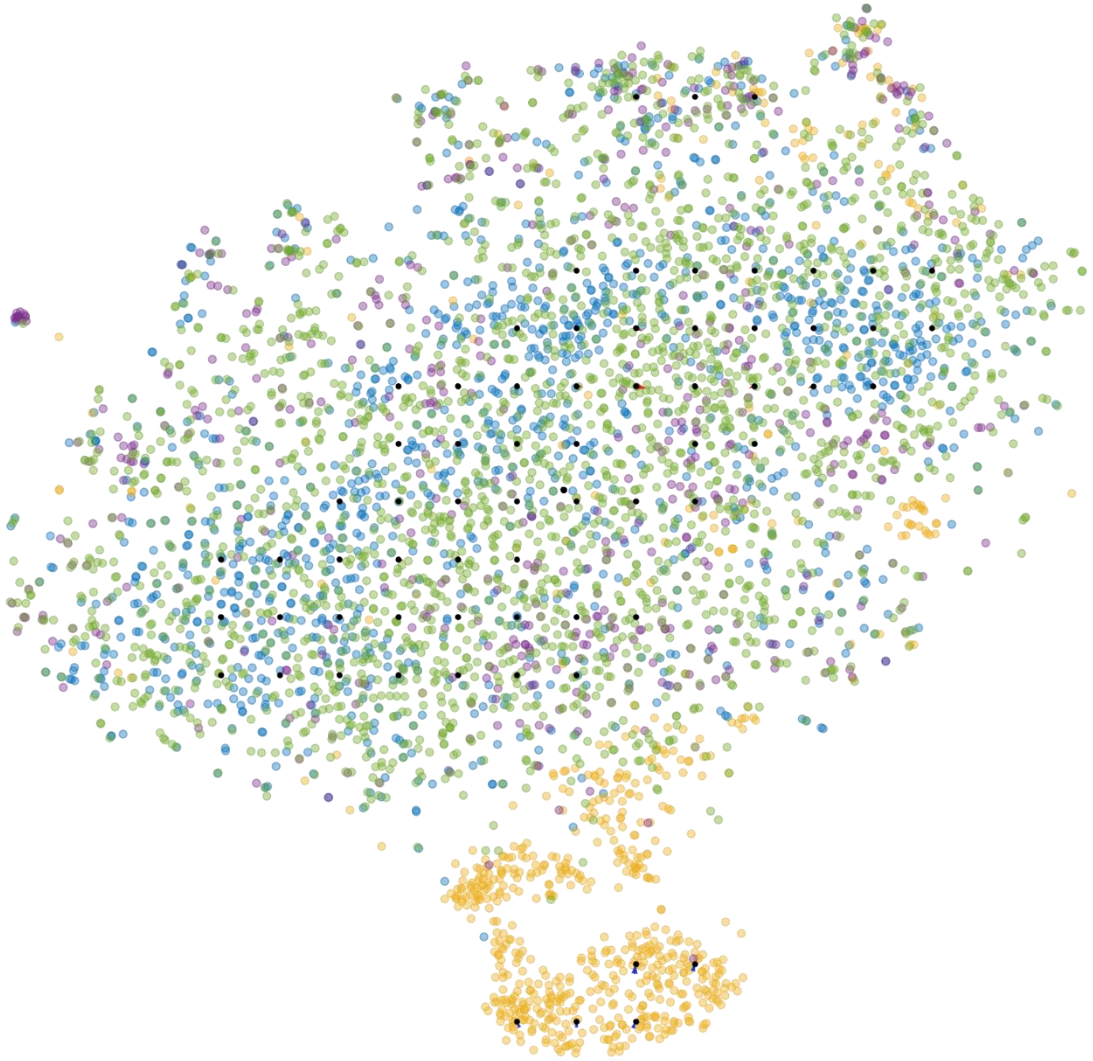

**Supplementary Figure 19.** ECCITE-seq ctrl velocity fields visualized on a grid. Arrow color identifies velocity estimate (RNA: red, protein: blue). Dot color identifies cell type (blue: CD4+ T, yellow: monocytes, green: CD8+ T, purple: natural killer).

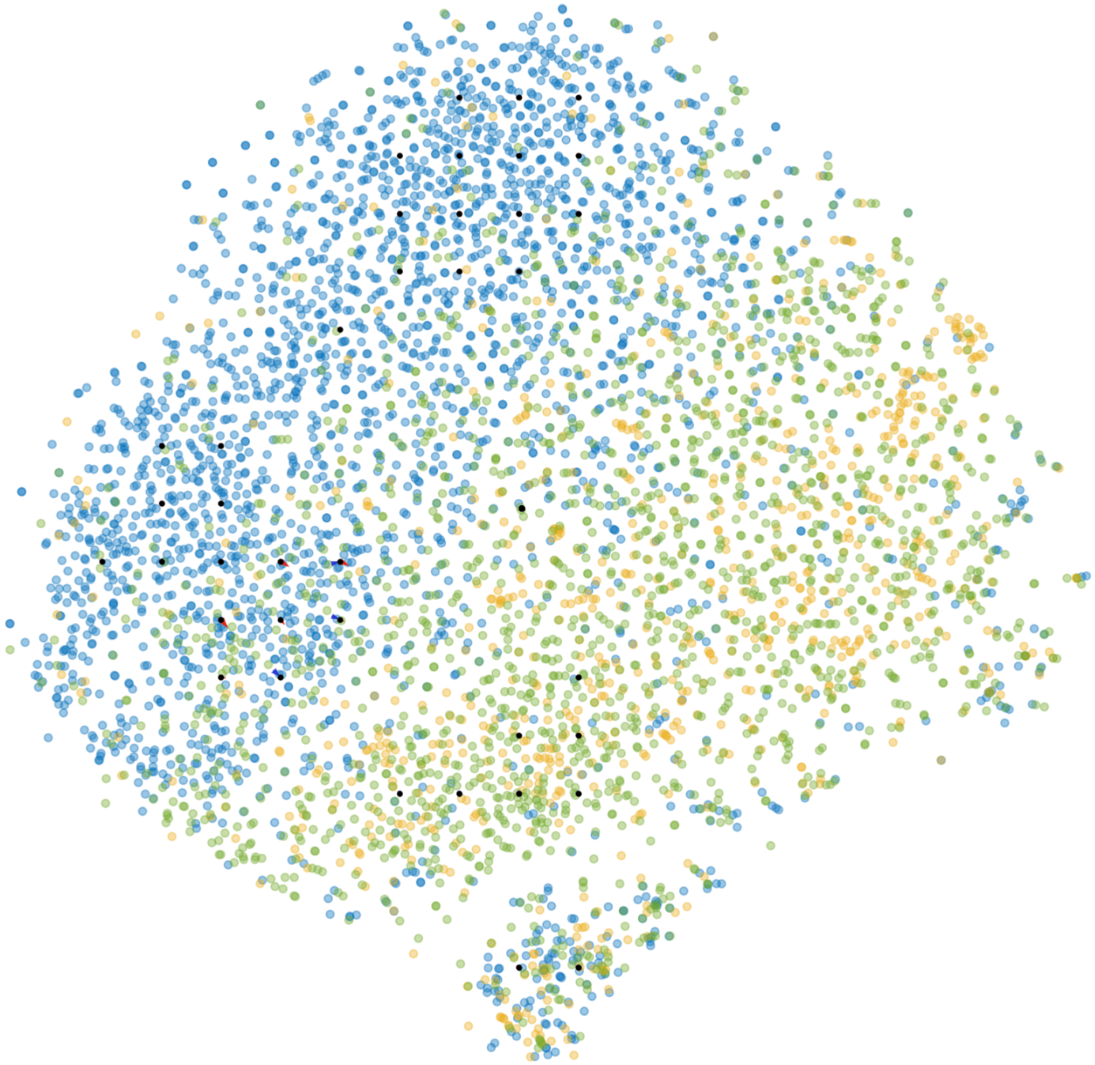

**Supplementary Figure 20.** ECCITE-seq CTCL velocity fields visualized on a grid. Arrow color identifies velocity estimate (RNA: red, protein: blue). Dot color identifies cell type (blue: CD4+ T, yellow: monocytes, green: CD8+ T).

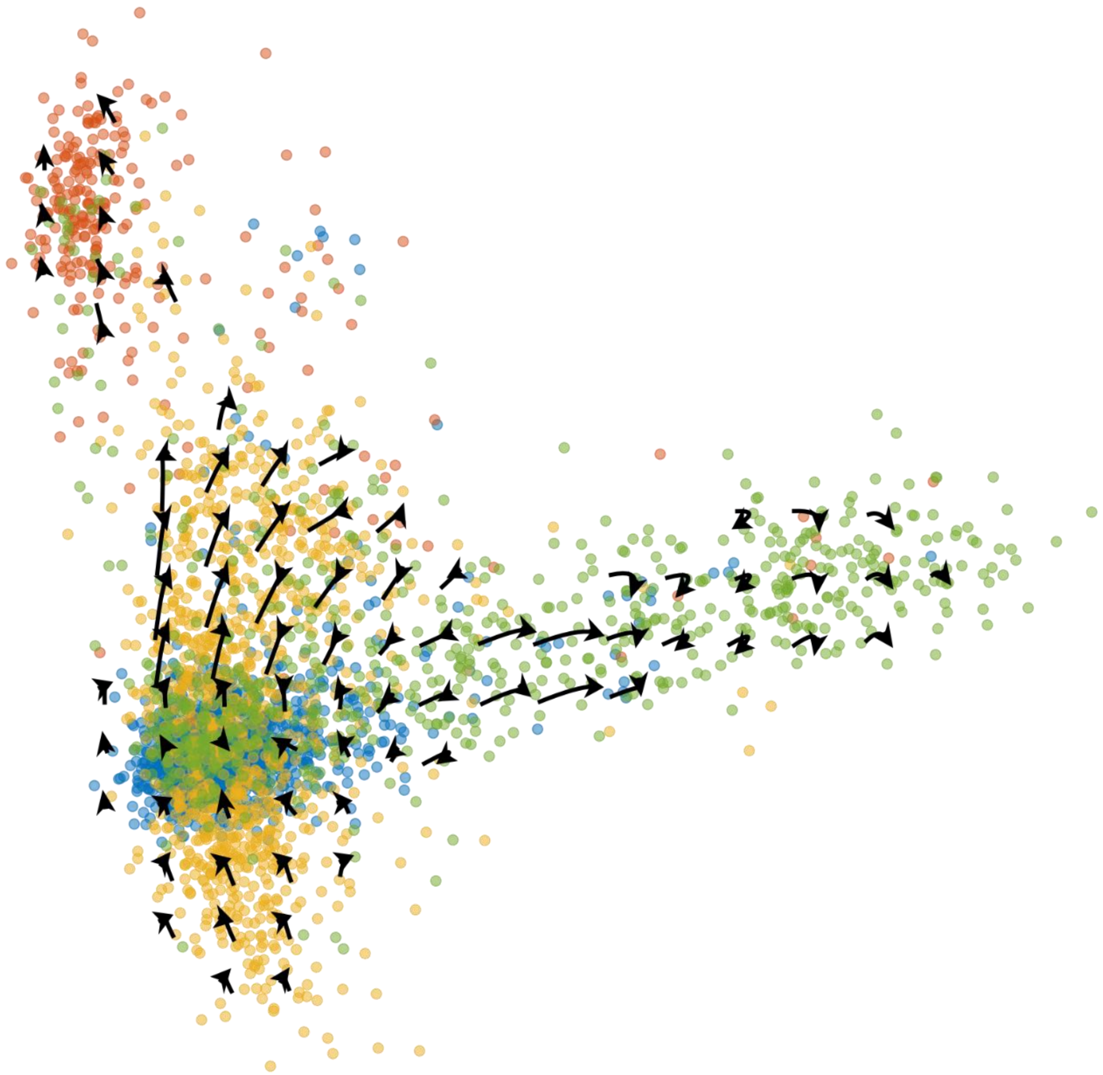

**Supplementary Figure 21.** REAP-seq combined velocity field. Dot color identifies cell type (blue: CD4+ T, red: B, yellow: monocytes, green: CD8+ T).

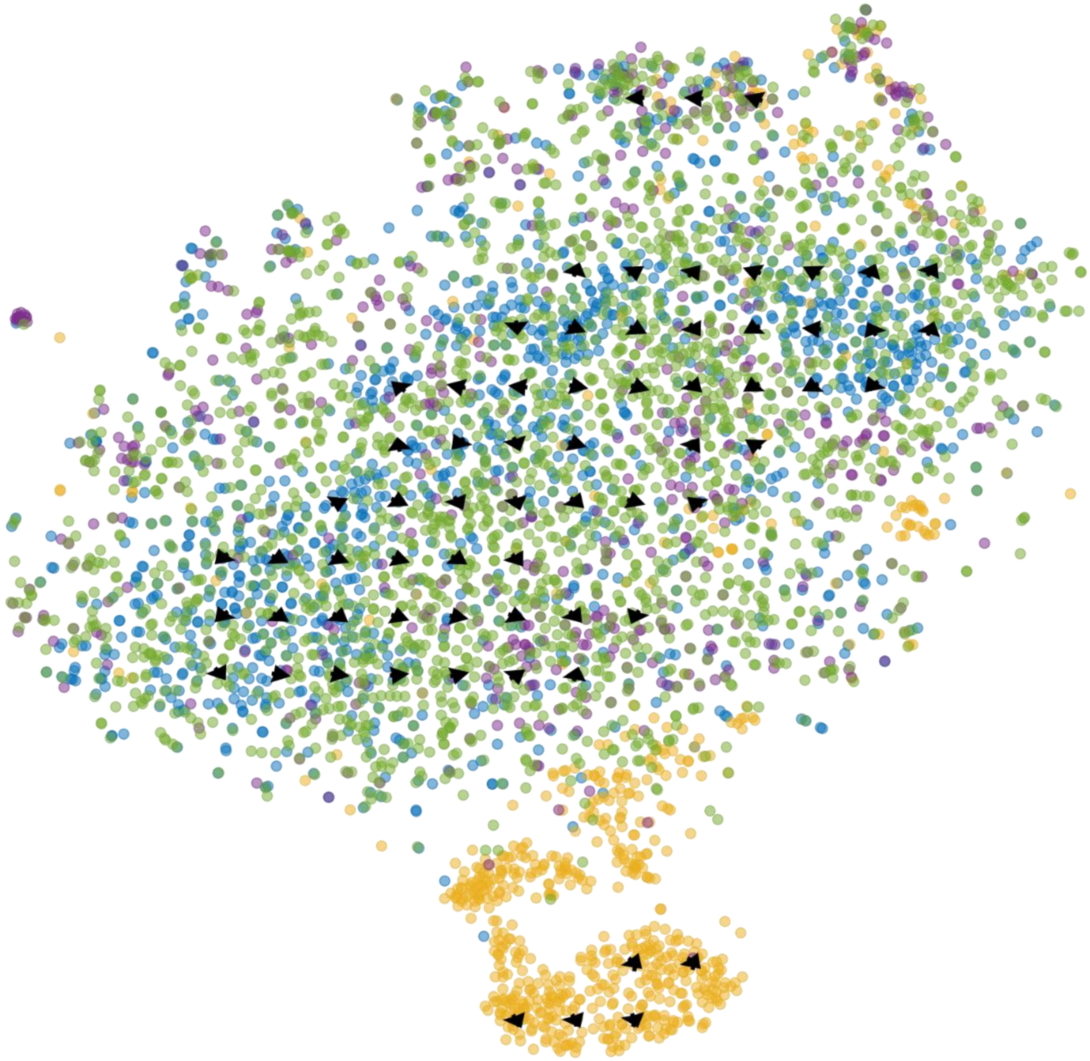

**Supplementary Figure 22.** ECCITE-seq ctrl combined velocity field. Dot color identifies cell type (blue: CD4+ T, yellow: monocytes, green: CD8+ T, purple: natural killer).

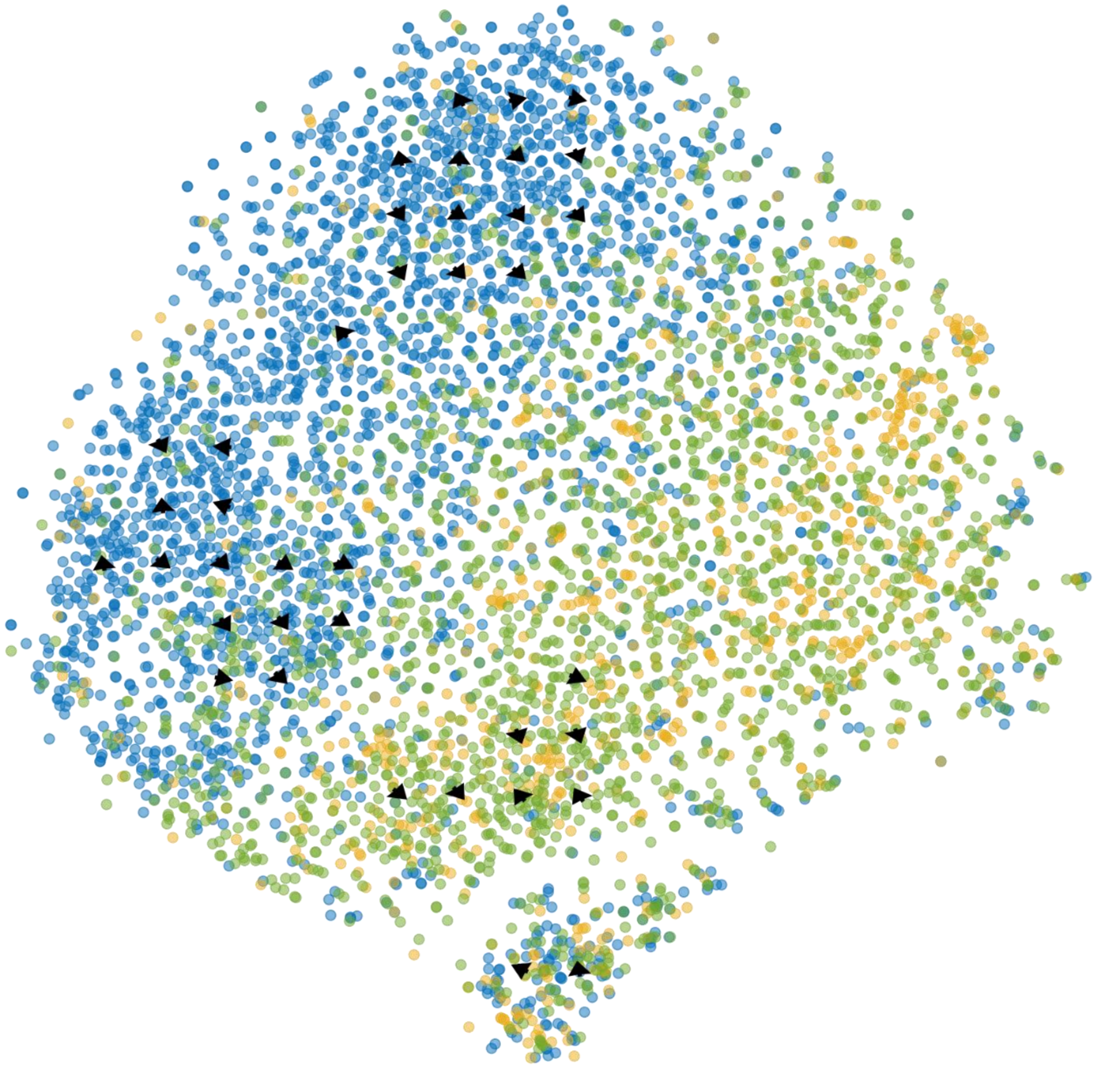

**Supplementary Figure 23.** ECCITE-seq CTCL combined velocity field. Dot color identifies cell type (blue: CD4+ T, yellow: monocytes, green: CD8+ T).

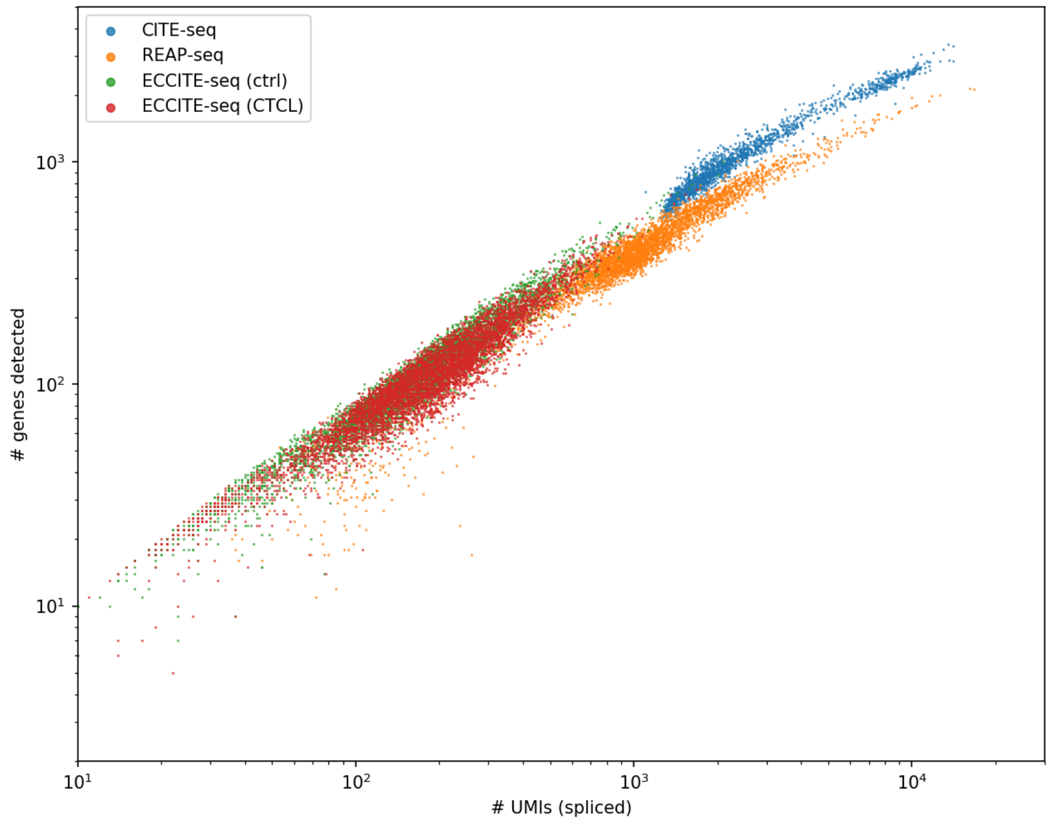


**Supplementary Figure 24**. Depth comparison between analyzed datasets. Each dot represents one cell in the dataset. The cell's number of spliced RNA molecules identified by the *velocyto* pipeline is shown on the abscissa; the number of genes with non-zero spliced RNA counts is shown on the ordinate.

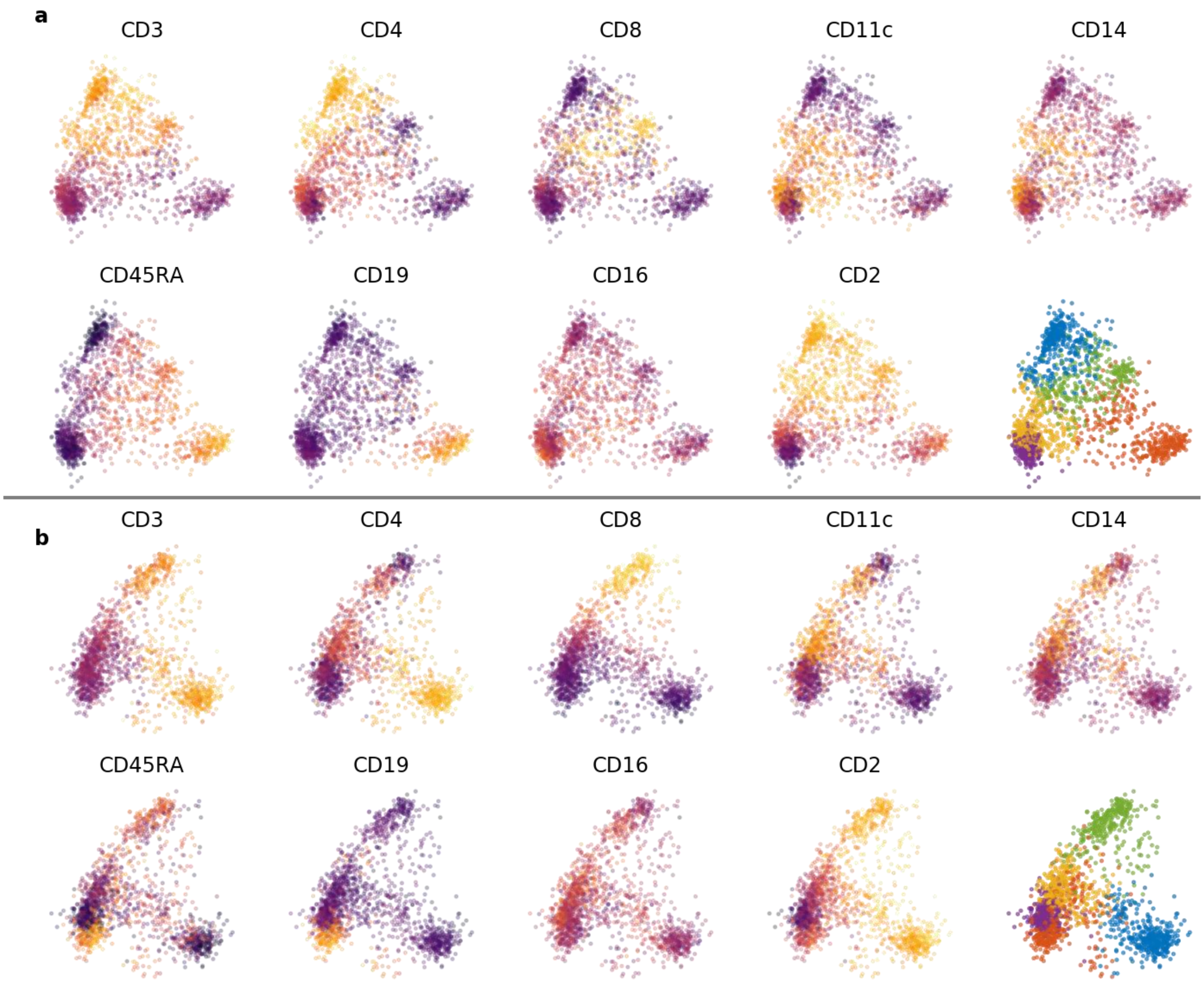


**Supplementary Figure 25.** CITE-seq cell cluster identification. (a) CITE-seq surface marker abundance (yellow is high) and cluster definitions; embedded in protein abundance PC1-PC2. (b) CITE-seq surface marker abundance and cluster definitions; embedded in protein abundance PC2-PC3. Color identifies cell type (blue: CD4+ T, red: B, yellow: monocytes, green: CD8+ T, purple: natural killer).

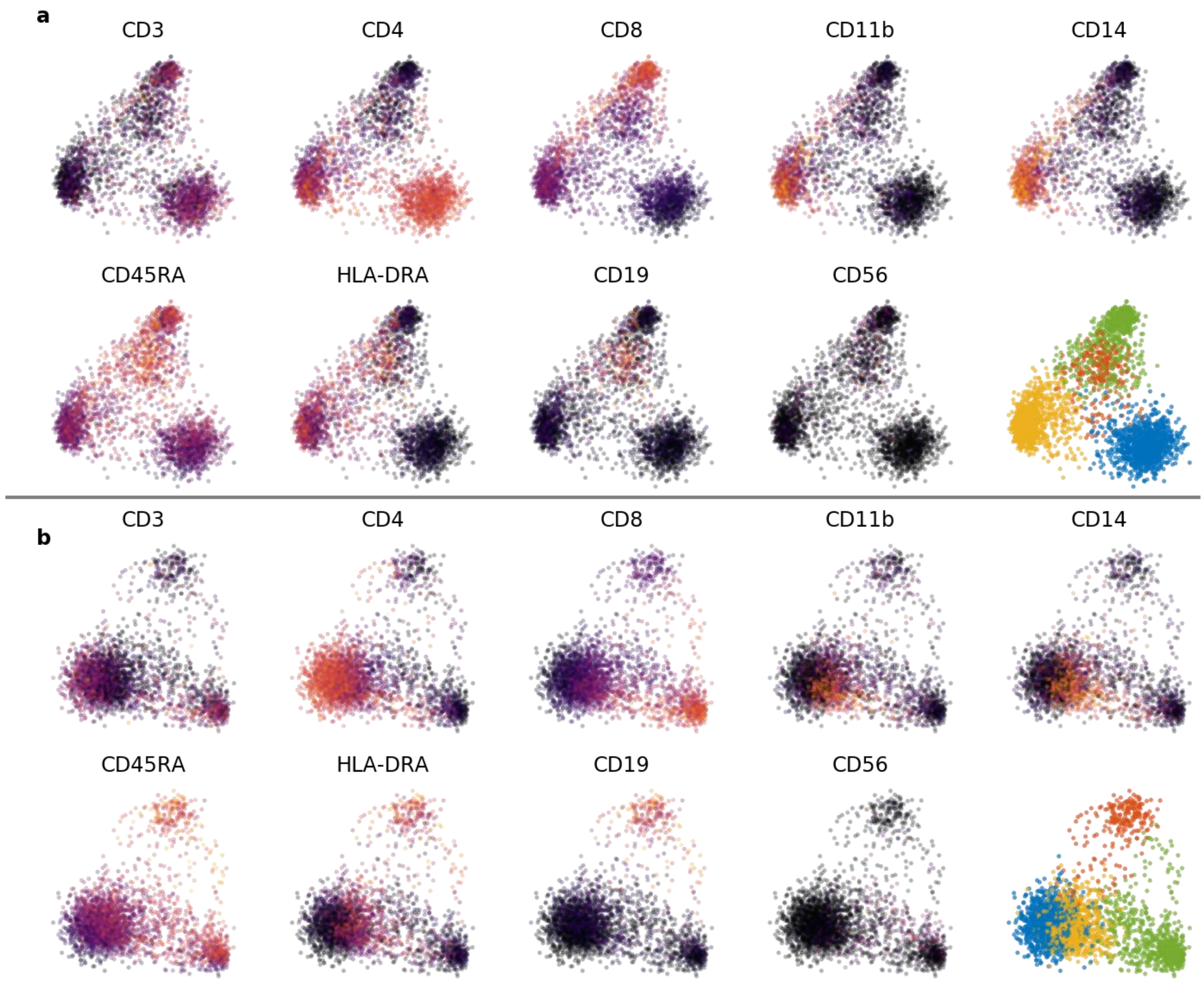


**Supplementary Figure 26.** REAP-seq cell cluster identification. (a) REAP-seq surface marker abundance (yellow is high) and cluster definitions; embedded in protein abundance PC1-PC2. (b) CITE-seq surface marker abundance and cluster definitions; embedded in protein abundance PC2-PC3. Color identifies cell type (blue: CD4+ T, red: B, yellow: monocytes, green: CD8+ T).

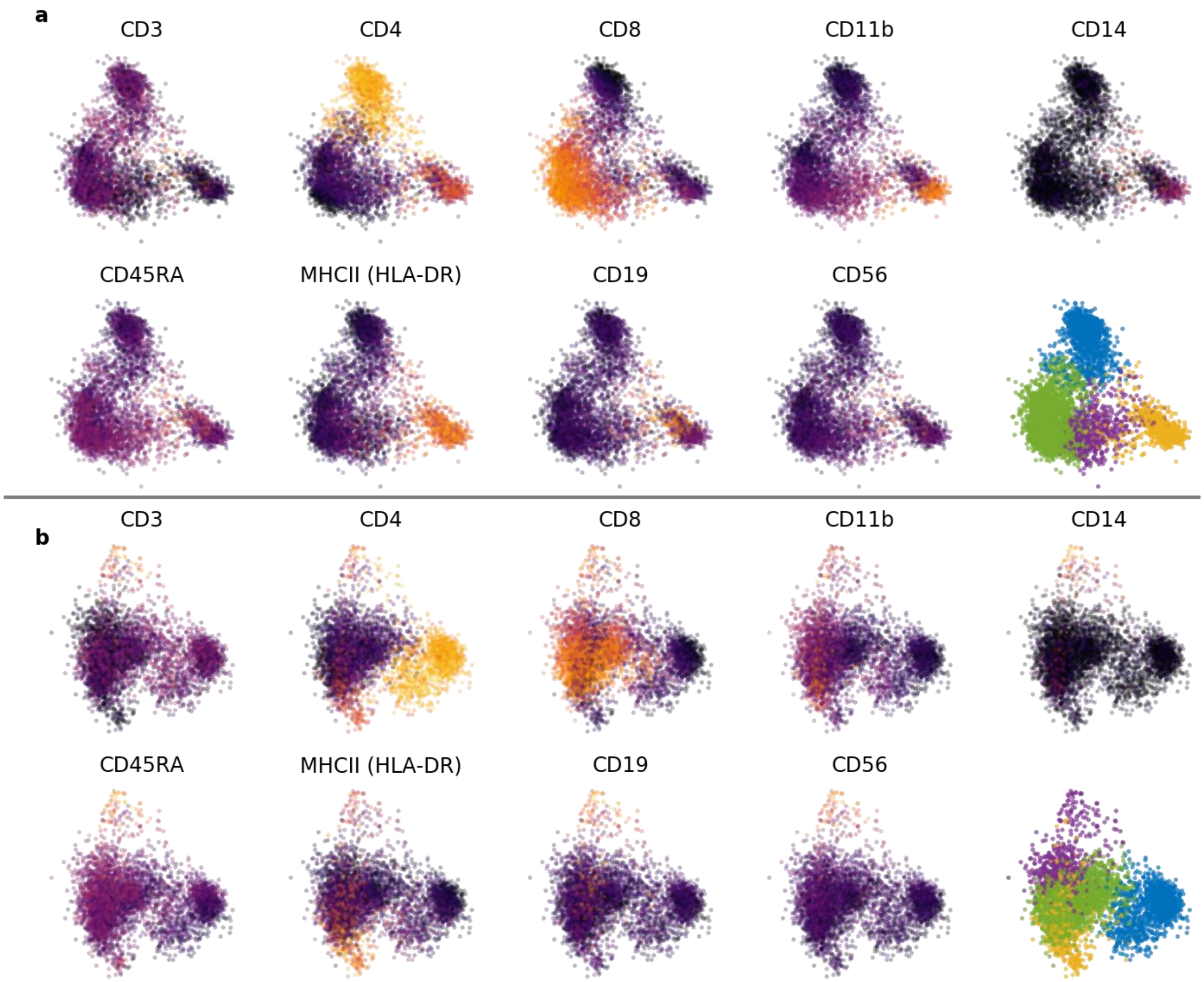


**Supplementary Figure 27.** ECCITE-seq ctrl cell cluster identification. (a) ECCITE-seq ctrl surface marker abundance (yellow is high) and cluster definitions; embedded in protein abundance PC1-PC2. (b) ECCITE-seq ctrl surface marker abundance and cluster definitions; embedded in protein abundance PC2-PC3. Color identifies cell type (blue: CD4+ T, yellow: monocytes, green: CD8+ T, purple: natural killer).

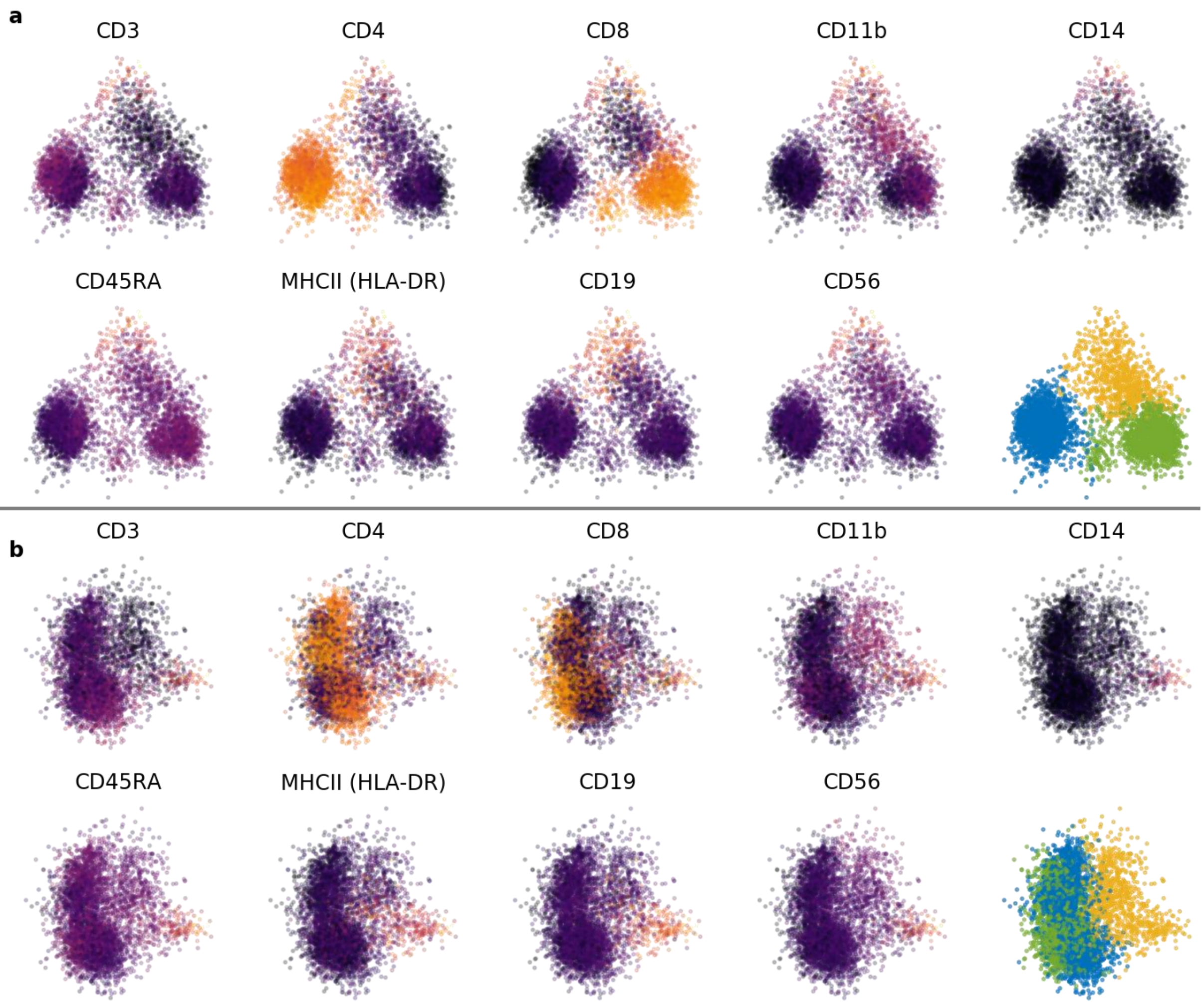


**Supplementary Figure 28.** ECCITE-seq CTCL cell cluster identification. (a) ECCITE-seq CTCL surface marker abundance (yellow is high) and cluster definitions; embedded in protein abundance PC1-PC2. (b) ECCITE-seq CTCL surface marker abundance and cluster definitions; embedded in protein abundance PC2-PC3. Color identifies cell type (blue: CD4+ T, yellow: monocytes, green: CD8+ T).
